# CUB domain-containing protein 1 controls HGF responses by integrating Src and Met–STAT3 signaling

**DOI:** 10.1101/789339

**Authors:** Kentaro Kajiwara, Atsuya Sugihara, Kazuhiro Aoki, Daisuke Okuzaki, Kunio Matsumoto, Masato Okada

**Affiliations:** Department of Oncogene Research, Research Institute for Microbial Diseases, Osaka University, Osaka 565-0871, Japan; Division of Quantitative Biology, Okazaki Institute for Integrative Bioscience, National Institute for Basic Biology, National Institutes of Natural Sciences, Aichi 444-8787, Japan; Genome Information Research Center, Research Institute for Microbial Diseases, Osaka University, Osaka 565-0871, Japan; Division of Tumor Dynamics and Regulation, Cancer Research Institute, Kanazawa University, Ishikawa 920-1192, Japan

## Abstract

Hepatocyte growth factor (HGF) controls various biological responses, including morphogenesis, organ regeneration, and cancer invasion, by activating its receptor, Met. However, the mechanisms that precisely control diverse Met signaling remain unclear. Here, we identified CUB domain-containing protein 1 (CDCP1) as a crucial element of HGF signaling. In MDCK cysts, HGF induced Src activation via CDCP1 upregulation, and CDCP1 ablation abrogated HGF responses, i.e., extended invasive cell protrusions and promoted cell growth/proliferation. Mechanistically, CDCP1 upregulation promoted Src recruitment into lipid rafts to activate Met-associated STAT3. In breast cancer cells in which Met and CDCP1 were co-upregulated, CDCP1 knockdown suppressed HGF-induced cell invasion. Furthermore, *in vivo* analysis showed that CDCP1 ablation suppressed compensatory renal growth by attenuating Met**–**STAT3 signaling. These findings demonstrate that CDCP1 plays a crucial role in controlling HGF responses by integrating Src and Met-STAT3 signaling, and provide new insights into the regulatory mechanisms underlying the multifaceted functions of HGF.

## Introduction

Hepatocyte growth factor (HGF) is a cytokine that was originally identified as a mitogenic factor for hepatocytes (Nakamura et al., 1989) and scatter factor (Weidner, Behrens, Vandekerckhove, & Birchmeier, 1990). HGF is secreted mainly from mesenchymal cells and acts on epithelial cells that express its receptor Met to control epithelial–mesenchymal interactions. Met was first isolated as an oncoprotein produced by a chromosomal translocation (Cooper et al., 1984) and was thereafter identified as a tyrosine kinase receptor for HGF (Bottaro et al., 1991). Met activation by HGF induces Met dimerization to activate autophosphorylation, resulting in the recruitment of multiple adaptor proteins (e.g., GAB1, GRB2 and SHC) and signal transducers (e.g., SOS, PI3K, and PLCγ-1), which subsequently activate downstream signaling pathways including the ERK, PI3K/Akt, NF-kB and STAT3/5 pathways (Trusolino, Bertotti, & Comoglio, 2010). Through activation of these multiple and complex signaling pathways, HGF–Met signaling controls diverse physiological responses, including developmental morphogenesis, tissue regeneration, and organ homeostasis (Nakamura, Sakai, Nakamura, & Matsumoto, 2011). Particularly, HGF plays a crucial role in epithelial tubulogenesis (Montesano, Matsumoto, Nakamura, & Orci, 1991), as well as dynamic cell migration and survival (Bladt, Riethmacher, Isenmann, Aguzzi, & Birchmeier, 1995).

The activation of HGF–Met signaling induces complex cellular responses that drive so-called “invasive growth,” including morphological changes associated with the epithelial–mesenchymal transition (EMT) and promotion of cell growth/proliferation (Gao & Vande Woude, 2005). The HGF–Met pathway is aberrantly upregulated in various cancers (Matsumoto, Umitsu, De Silva, Roy, & Bottaro, 2017), such as breast, esophageal, hepatocellular carcinoma, and non-small cell lung cancers. The invasive-growth phenotypes induced by HGF contribute to the progression of invasion and metastasis, as well as the growth/survival of cancer cells. Thus, HGF–Met signaling has been considered as a promising therapeutic target for a subset of cancers (Comoglio, Trusolino, & Boccaccio, 2018).

HGF also plays a role in the compensatory growth of organs after loss of their mass and/or function (Trusolino et al., 2010). Upon unilateral nephrectomy (UNX), HGF is produced by the surrounding or distal mesenchyme in the remaining kidney (Ishibashi et al., 1992; Joannidis, Spokes, Nakamura, Faletto, & Cantley, 1994; Nagaike et al., 1991) and induces the Met upregulation in the renal tubules (Ishibashi et al., 1992). HGF facilitates dynamic morphogenesis through the induction of EMT and cell growth in renal epithelial cells (Chang-Panesso & Humphreys, 2017; Matsumoto & Nakamura, 2001). This compensatory renal growth is promoted via the activation of mTOR (Chen, Chen, Neilson, & Harris, 2005; Chen, Chen, Thomas, Kozma, & Harris, 2009; Chen et al., 2015) and STAT3 signaling (Boccaccio et al., 1998; Kermorgant & Parker, 2008). However, the exact molecular mechanisms controlling HGF-Met signaling during compensatory organ growth remain elusive.

The Madin–Darby canine kidney (MDCK) cell line was derived from renal tubule epithelial cells and is a physiologically relevant in vitro model for studying the regulation of HGF functions (O’Brien, Zegers, & Mostov, 2002; Zegers, 2014). When grown in three-dimensional cultures, MDCK cells spontaneously form spherical cysts that resemble renal tubules, comprising an epithelial monolayer and lumen. Upon stimulation with HGF, MDCK cysts undergo morphological alterations and form branched tubular structures (Montesano, Matsumoto, et al., 1991; Montesano, Schaller, & Orci, 1991). During this morphogenesis, MDCK cells lose their epithelial polarity via a partial EMT-like phenotypic change and protrude into the extracellular matrix (ECM) by degrading the basement membrane (O’Brien et al., 2004; Pollack, Runyan, & Mostov, 1998). In addition, HGF promotes cell growth and proliferation, resulting in the formation of multi-cell layered cysts. These observations indicate that HGF can induce invasive growth in MDCK cysts. However, how multiple signaling pathways downstream of Met are controled to properly execute HGF-induced invasive growth responses has not been addressed.

To address the above issues, we tested the hypothesis that there an additional factor(s) that selectively links and/or controls critical pathways required for HGF-induced invasive growth responses. To this end, we re-dissected the HGF–Met signaling pathway using three-dimensional cultures of MDCK cysts and identified Src and its membrane-scaffold protein, CUB domain-containing protein 1 (CDCP1), as critical elements of HGF–Met signaling. We then investigated the mechanisms underlying the functions of CDCP1 in MDCK cysts, and confirmed the role of CDCP1 in HGF-induced cancer migration and invasion, using breast cancer MDA-MB231 cells with increased Met and CDCP1 expression. Finally, we verified *in vivo* roles of CDCP1 in compensatory renal growth using *Cdcp1*-knockout mice. The results show that CDCP1 plays a crucial role in controlling the HGF-induced invasive growth responses by focally and temporally integrating Src and Met–STAT3 signaling in lipid rafts.

## Results

### CDCP1 is required for HGF signaling in MDCK cysts

MDCK cells cultured in a collagen matrix formed cyst structures with a luminal space (Fig. 1A). HGF treatment immediately promoted cell growth/proliferation (as indicated by increased cyst diameters) and morphological changes with some cysts exhibiting extended cell protrusions (Fig. 1, A–C). To dissect the HGF-induced signaling pathways, we first examined the effects of various signaling inhibitors on HGF-induced phenomena. Treatment of MDCK cysts with NK4, a specific HGF antagonist (Date, Matsumoto, Shimura, Tanaka, & Nakamura, 1997), robustly inhibited cell proliferation and morphological changes, confirming that the observed phenomena were dependent on HGF. Treatment with the selective mTOR inhibitors, Torin1 and rapamycin, significantly suppressed cell growth/proliferation but was less effective against the formation of cell protrusions (Fig. 1, A–C). In contrast, treatment with dasatinib (a Src kinase inhibitor) potently suppressed cell proliferation and protrusion extension (Fig. 1, A–C). Furthermore, immunofluorescence analysis revealed that activated Src (pY418) was concentrated at the tip of protruding cells (Fig. 1D), which was visualized with mCherry-GPI, a marker protein of lipid rafts (Fig. 1E). These observations imply that HGF signaling is associated with Src activation in lipid rafts on protruding cells.

**Fig. 1.**
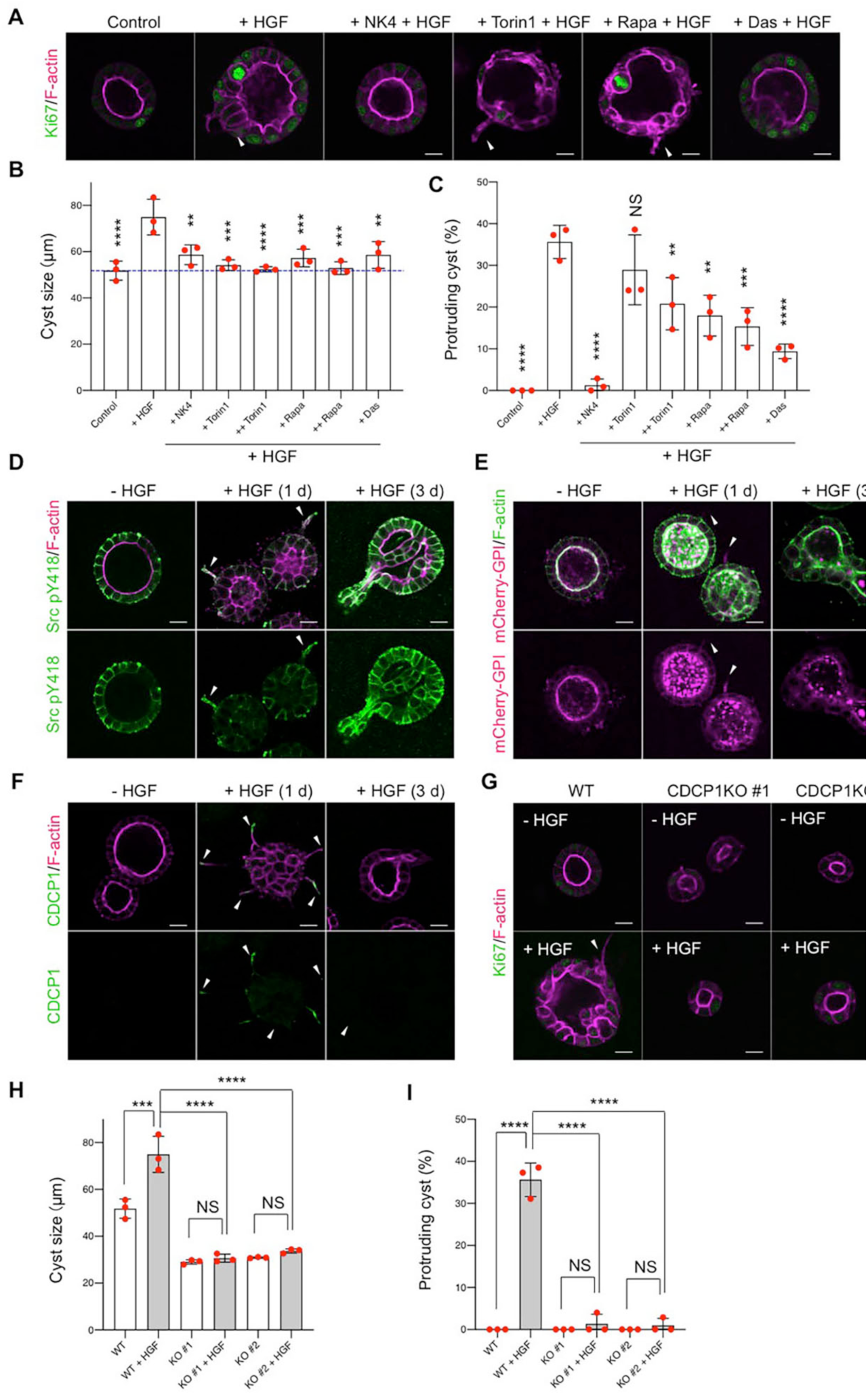
CDCP1 is required for HGF-induced invasive growth. (**A**) MDCK cysts embedded within the collagen matrix were pretreated with NK4 (1 µg/ml), Torin1 (+, 50 nM; ++, 100 nM), rapamycin (+ Rapa, 50 nM; ++ Rapa, 100 nM), or dasatinib (20 nM, Das), and then incubated in the presence of HGF (50 ng/ml) for one day. Ki67 was visualized with an Alexa Fluor 488-conjugated antibody (green), and actin filaments were stained with Alexa Fluor 594-phalloidin (magenta). The arrowheads indicate transiently formed protrusions. (**B**) Diameter (µM) of cysts (n = 100). The dotted blue line indicates the average diameter of non-treated cysts. (**C**) Fraction of the total number of cysts counted (n > 100) with protrusions. (**D**) MDCK cysts embedded within the collagen matrix were incubated in the presence of HGF for the indicated time periods. Activated Src (pY418) was visualized with an Alexa Fluor 488-conjugated antibody (green), and actin filaments were stained with Alexa Fluor 594-phalloidin (magenta). The arrowheads indicate transiently formed protrusions. (**E**) mCherry-GPI-overexpressing MDCK cysts embedded within the collagen matrix were incubated in the presence of HGF for the indicated time periods. Actin filaments were stained with Alexa Fluor 488-phalloidin (green). (**F**) MDCK cysts embedded within the collagen matrix were incubated in the presence of HGF for the indicated time periods. The localization of CDCP1 was visualized using an Alexa Fluor 488-conjugated antibody (green), and actin filaments were stained with Alexa Fluor 594-phalloidin (magenta). (**G**) Wild-type and CDCP1-knockout MDCK cysts were incubated in the presence of HGF for 1 day. Ki67 was visualized with an Alexa Fluor 594-conjugated antibody (magenta), and actin filaments were stained with Alexa Fluor 488-phalloidin. The arrowheads indicate transiently formed protrusions. The scale bars indicate 50 µm. (**H**) Diameter (µm) of cysts (n = 100). (**I**) Fraction of the total number of cysts counted (n > 100) with protrusions. The scale bars indicate 10 µm. The mean ratios ± SD were obtained from three independent experiments. **, *P* < 0.01; ***, *P* < 0.001; ****, *P* < 0.0001; NS, not significantly different; ANOVA was calculated compared to HGF-treated cysts.

To assess the role of Src in HGF signaling, we examined the effects of Src activation on MDCK cysts by expressing a Src protein fused to a modified estrogen receptor (Src–MER) (Kuroiwa, Oneyama, Nada, & Okada, 2011) that could be activated by treatment with hydroxytamoxifen (4-OHT; Fig. S1A). Src activation induced both the formation of multiple cell protrusions and cell proliferation (Fig. S1B). Detergent-resistant membrane (DRM)-separation analysis revealed that activated Src–MER accumulated in lipid raft fractions (Fig. S1C). We thus searched for scaffolding proteins that accommodate and activate Src in these fractions. Src–MER-interacting proteins were isolated from DRM fractions by co-immunoprecipitation and were identified by mass spectrometry (Fig. S1, D and E). Among the candidate proteins identified, we selected the transmembrane glycoprotein CDCP1 for further analysis because it has palmitoylation sites required for lipid raft localization and serves as a membrane scaffold of Src (Alvares, Dunn, Brown, Wayner, & Carter, 2008; Wortmann, He, Deryugina, Quigley, & Hooper, 2009) (Fig. S2A). Immunofluorescence analysis revealed that endogenous CDCP1 was transiently concentrated at the tips of protruding cells during morphological changes (Fig. 1F), in a manner similar to that of activated Src and mCherry–GPI (Fig. 1, D and E). More importantly, HGF-induced invasive growth phenotypes were efficiently abrogated by *Cdcp1*-knockout in MDCK cells (Fig. 1, G–I and Fig. S2, B–D). These results suggest that the CDCP1 is a crucial component of HGF signaling.

### CDCP1 upregulation phenocopies HGF-induced invasive growth by activating Src in lipid rafts

To elucidate the mechanisms whereby CDCP1 regulates HGF signaling, we generated MDCK cells with doxycycline (Dox)-inducible in CDCP1 expression. Dox treatment induced the expression of full-length and cleaved CDCP1 at the plasma membrane (Fig. S3, A and B). DRM-separation analysis showed that a fraction of CDCP1 was distributed in lipid raft fractions (Fig. S3C). Moreover, the induction of CDCP1 expression dramatically activated Src (pY418/Src) (Fig. S3A). Furthermore, activated Src was concentrated in the lipid raft fractions and was associated with CDCP1 (Fig. S3, C and D). In contrast, a CDCP1 mutant that lacked the Src-binding site (Y734F; CDCP1-YF) failed to activate Src (Fig. S3A). A different CDCP1 mutant lacking the lipid raft localization signal (residues C689G and C690G; CDCP1-CG) could activate Src, but only in non-lipid raft fractions (Fig. S3, A and C). These findings indicate that upregulated CDCP1 activates Src in lipid rafts.

Next, we examined the effects of CDCP1 expression on the phenotypes of MDCK cysts (Fig. 2A). CDCP1 expression was induced after the completion of cystogenesis and morphological changes were observed (Movie S1). In the early stages (12–24 h after induction), cells expressing CDCP1 gradually protruded toward the ECM. During the later stages (48 h post-induction), multiple cell protrusions formed randomly and extended aggressively toward the ECM, with cysts forming multiple layers (Fig. 2B). Concurrent with these phenomena, CDCP1 upregulation activated Src in the protruding cells (Fig. 2C) and Src inhibition by dasatinib strongly suppressed CDCP1-induced cell proliferation and protrusion extension (Fig. 2D and Table S1). In addition, the upregulation of CDCP1-YF or CDCP1-CG did not significantly induce phenotypic changes (Fig. 2, E and F). Based on these results, it is likely that upregulation of CDCP1 can phenocopy HGF-induced invasive growth phenotypes via Src activation in lipid rafts.

**Fig. 2.**
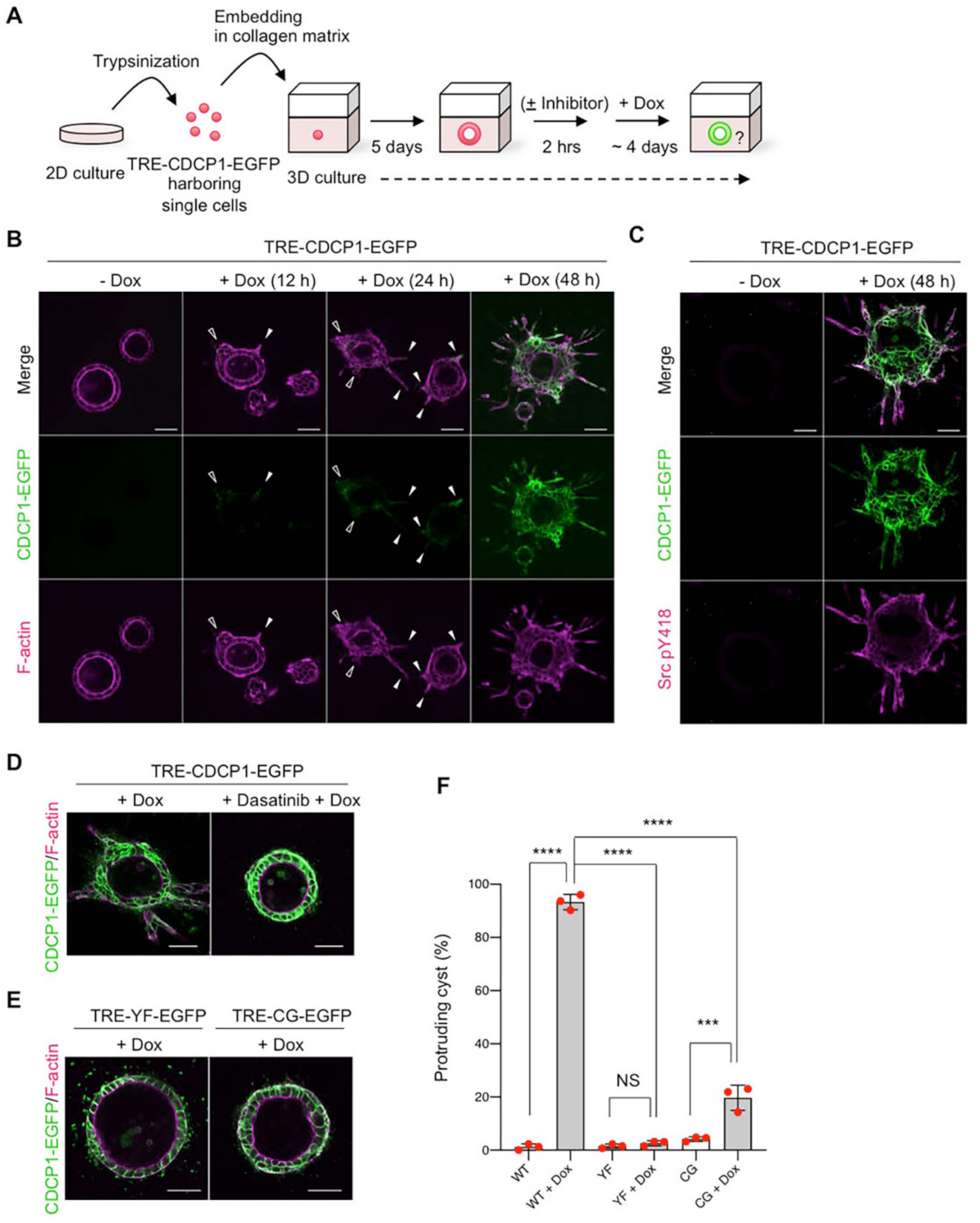
CDCP1 upregulation phenocopies HGF-induced invasive growth by activating Src in lipid rafts. (**A**) Schematic representation of the procedure used to analyze TRE–CDCP1–EGFP-harboring MDCK cysts (CDCP1–EGFP cysts). TRE–CDCP1–EGFP-harboring cells were cultured within a collagen matrix for 5 days to allow time for cyst formation, and then CDCP1–EGFP cysts were incubated in the presence of Dox for ∼4 days. (**B**) CDCP1–EGFP cysts embedded within the collagen matrix were incubated in the presence of Dox (1 µg/ml) for the indicated time periods. Actin filaments were stained with Alexa Fluor 594-phalloidin (magenta). Filled arrowheads indicate protruding cells and open arrowheads indicate multi-layered structures. (**C**) CDCP1– EGFP cysts were incubated in the presence of Dox for 2 days. Activated Src was visualized using an anti-Src pY418 antibody (magenta). (**D**) CDCP1–EGFP cysts embedded within the collagen matrix were pretreated with 20 nM dasatinib for 2 h, and then incubated with Dox for 4 days. (**E**) CDCP1–YF-EGFP and CDCP1–CG-EGFP cysts embedded within the collagen matrix were incubated in the presence of Dox for 4 days. Actin filaments were stained with Alexa Fluor 594-phalloidin (magenta). The scale bars indicate 50 µm. (**F**) Fraction of the total number of cysts counted (n > 150) with protrusions. The mean ratios ± SDs were determined from three independent experiments. ***, *P* < 0.001; ****, *P* < 0.0001; NS, not significantly different; ANOVA compared to the Dox-treated cysts.

### CDCP1 activates the STAT3 pathway via lipid rafts

To dissect the signaling pathways operating downstream of the CDCP1–Src axis, we performed DNA-microarray analyses of MDCK cysts expressing either wild-type CDCP1 or the CDCP1-CG mutant. Gene ontology analysis implied that CDCP1 expression participated in regulating signal transduction (Fig. S4A) and Ingenuity Pathway Analysis revealed that the STAT3 pathway was more prominently activated by CDCP1 expression than by CDCP1-CG expression (Fig. S4, B and C, Data file S1). Indeed, the level of phosphorylated STAT3 (pY705) increased significantly in cysts expressing wild-type CDCP1 (Fig. 3A). STAT3 is a transcription factor that is activated by Src-mediated phosphorylation of Tyr705 (Bromberg, Horvath, Besser, Lathem, & Darnell, 1998) and regulates various cellular events, including the upregulation of matrix metalloproteinase (MMPs) (H. Yu, Pardoll, & Jove, 2009) and mitogenic genes.

**Fig. 3.**
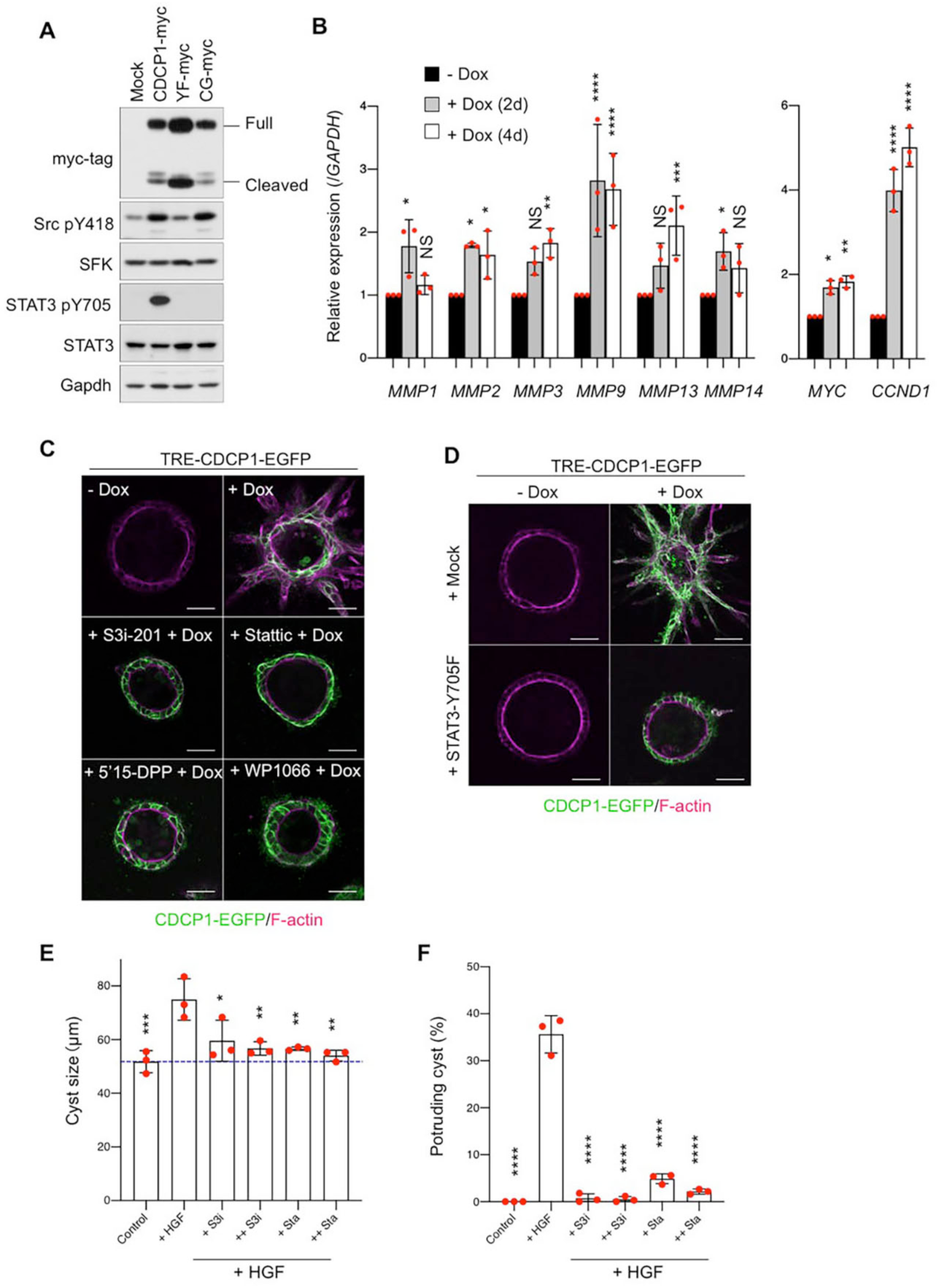
STAT3 activation is required for CDCP1-induced invasive growth. (**A**) MDCK cells overexpressing wild-type or mutant CDCP1 (Myc-tagged variants) were embedded within the collagen matrix and cultured for 9 days. Cyst lysates were subjected to immunoblotting using the indicated antibodies. (**B**) CDCP1–EGFP cysts were incubated with Dox (1 µg/ml) for 2 or 4 days, and then subjected to quantitative real-time PCR. Relative mRNA-expression levels were calculated by setting the mean value for non-treated cysts to 1. The mean ratios ± SDs were determined from three independent experiments. ANOVA calculations were performed to compare the results with those from non-treated cysts. (**C**) CDCP1–EGFP cysts were pretreated with the indicated STAT3-specific inhibitors for 2 h and then incubated with Dox (1 µg/ml) for 4 days.. (**D**) STAT3–Y705F-overexpressing CDCP1–EGFP cysts were incubated with Dox (1 µg/ml) for 4 days. Actin filaments were stained with Alexa Fluor 594-phalloidin (magenta). The scale bars indicate 50 µm. (**E, F**) MDCK cysts embedded within the collagen matrix were pretreated with S3i-201 (+ S3i, 50 nM; ++ S3i, 100 nM) or Stattic (+ Sta, 2.5 nM; ++ Sta, 5.0 nM) for 2 h and then incubated with HGF (50 ng/ml) for 1 day. (**E**) Diameter (µm) of cysts (n = 100). (**F**) Fraction of the total number of cysts counted (n > 100) with protrusions. The mean ratios ± SDs were obtained from three independent experiments. *, *P* < 0.05; **, *P* < 0.01; ***, *P* < 0.001; ****, *P* < 0.0001; NS, not significantly different; ANOVA was calculated compared to the HGF-treated cysts.

In fact, the expression levels of several MMP-encoding genes were significantly increased by CDCP1 upregulation (Fig. 3B). In addition, immunofluorescent staining of laminin revealed that CDCP1 expression disrupted the basement membrane, enabling the extension of cell protrusions (Fig. S5A). This effect was suppressed by treatment with marimastat, a pan-MMP inhibitor (Fig. S5, B and C). Similar suppression of cell protrusion extension by marimastat was also observed in HGF-stimulated MDCK cysts (Fig. S5, D and E). Analysis with DQ-collagen, a fluorescent indicator of collagen degradation, also showed that CDCP1 expression induced ECM degradation (Fig. S5F). These observations suggest that the HGF-induced formation of cell protrusions is associated with ECM rearrangements via STAT3-mediated upregulation of MMPs. Mitogenic genes such as *Myc* (Myc) and *Ccnd1* (Cyclin D1) were also upregulated in cysts expressing CDCP1 (Fig. 3B), indicating that CDCP1-induced cell proliferation is attributable to STAT3 activation.

We further confirmed that STAT3 contributes to HGF signaling by specifically perturbing its activity. Activation of STAT3 using the STAT3–MER system induced the formation of cell protrusions and a multi-layered structure (Fig. S5G). In contrast, treatment of MDCK cysts with STAT3-specific inhibitors suppressed CDCP1-induced cell growth (Fig. 3C and Table S1), and overexpression of dominant negative STAT3 (STAT3-Y705F) also inhibited CDCP1-induced cellular events (Fig. 3D). Furthermore, STAT3 inhibitors efficiently suppressed cell proliferation (Fig. 3E) and protrusion formation (Fig. 3F) in HGF-stimulated MDCK cysts, underscoring the crucial role of STAT3 activation in HGF-induced invasive growth.

### CDCP1 focally integrates Src and Met-STAT3 signaling

Next, we investigated the functional link between the HGF–Met pathway and the CDCP1–Src axis. An inhibitor screen assay showed that various Met-specific inhibitors potently suppressed CDCP1-induced invasive growth (Fig. 4A and Table S1), suggesting that Met activity facilitated the function of CDCP1. Because activated Met associates with STAT3, we predicted a close association between the CDCP1–Src and Met–STAT3 axes. To assess physical interactions between these proteins, Met and various CDCP1 mutants that lacked particular extracellular CUB domains were co-expressed in HEK293 cells (Fig. 4, B and C). Co-immunoprecipitation assays revealed that CDCP1 interacted with Met. This interaction was enhanced by removing the first CUB domain of CDCP1, but was diminished by deleting all CUB domains (Fig. 4, B and C), indicating that CDCP1 interacts with Met through extracellular domains and that removal of the first CDCP1 CUB domain is required for efficient association with Met.

**Fig. 4.**
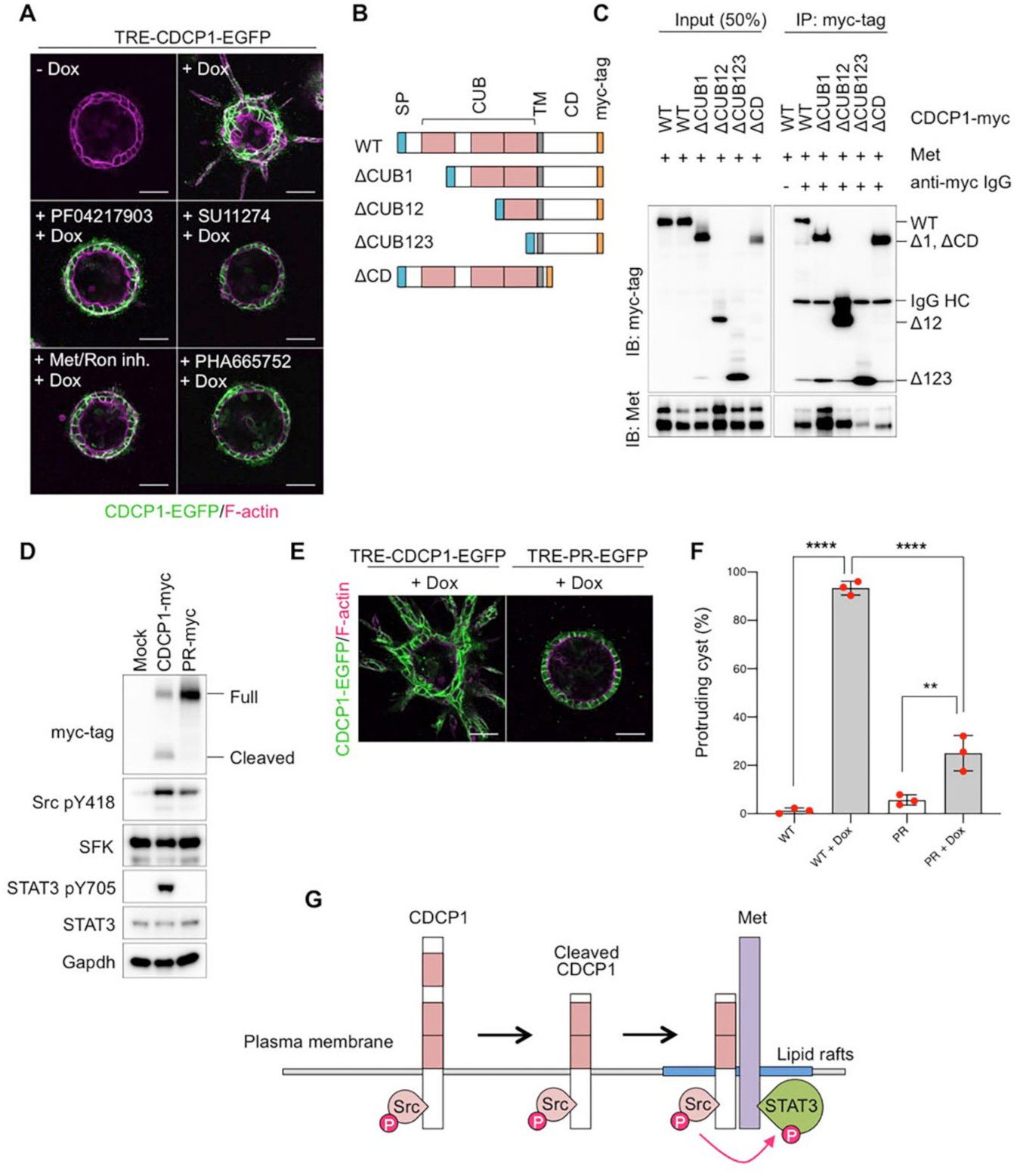
The CDCP1–Met association is required for CDCP1-induced invasive growth. (**A**) CDCP1–EGFP cysts were pretreated with the indicated Met-specific inhibitors for 2 h and then incubated with Dox (1 µg/ml) for 4 days. (**B**) Schematic representation of CDCP1 and the deletion mutants. SP, signal peptide; CUB, CUB domain; TM, transmembrane domain; CD, cytosolic domain. (**C**) Lysates from HEK293 cells overexpressing both CDCP1-Myc and Met were subjected to immunoprecipitation with an anti-Myc tag antibody. Immunoprecipitates were subjected to immunoblotting using the indicated antibodies (IgG HC, IgG heavy chain). (**D**) CDCP1-myc- and CDCP1-PR-myc-overexpressing MDCK cells were embedded within collagen matrix and cultured for 9 days. Cyst lysates were subjected to immunoblotting using the indicated antibodies. (**E**) CDCP1–PR–EGFP cysts embedded within the collagen matrix were incubated with Dox for 4 days. Actin filaments were stained with Alexa Fluor 594-phalloidin (magenta). The scale bars indicate 50 µm. (**F**) Fraction of the total number of cysts counted (n > 150) with protrusions. The mean ratios ± SDs were obtained from three independent experiments. **, *P* < 0.001; ****, *P* < 0.0001; NS, not significantly different; ANOVA was calculated compared to the Dox-treated cysts. (**G**) Schematic model of the role of CDCP1–Met association in Src-induced STAT3 phosphorylation.

Because CDCP1 is activated via proteolytic shedding between the first and second CUB domains (Casar et al., 2012; He, Harrington, & Hooper, 2016), it is likely that activated CDCP1 functionally interacts with Met. To test this hypothesis, we analyzed the function of a mutant CDCP1 (CDCP1-PR) rendered resistant to proteolytic shedding due to the presence of three point mutations (K365A, R368A, and K369A; Fig. 4D). CDCP1-PR expression induced Src activation to a level similar to that observed with wild-type CDCP1, but failed to activate STAT3 (Fig. 4D). Consistent with these biochemical effects, CDCP1-PR failed to induce the formation of cell protrusions (Fig. 4, E and F). These data suggest that an association between activated CDCP1 and Met is required for Src-mediated STAT3 activation and subsequent cellular events (Fig. 4G). In addition, the initiation of cell protrusion induced by HGF treatment or Met overexpression was efficiently suppressed by the loss of CDCP1 (Fig. S6A). These lines of in vitro evidence suggest a model wherein CDCP1–Src upregulation in lipid rafts is required for Met-mediated STAT3 activation during HGF-induced invasive growth in MDCK cysts (Fig. S6B).

### CDCP1 is required for HGF-induced cancer invasion

To further explore the role of the CDCP1–Met interaction in other HGF-dependent cellular responses, we examined whether CDCP1 was involved in HGF-induced invasion of cancer cells. CDCP1 is upregulated in various cancers (He et al., 2016), including malignant triple-negative breast cancers (Turdo et al., 2016). Indeed, highly invasive triple-negative breast cancer MDA-MB231 cells expresses CDCP1 at high levels (Fig. 5A), while non-invasive luminal A-type breast cancer MCF7 and T47D cells do not express CDCP1. Notably, Met is also upregulated in MDA-MB231 cells, but not in MCF7 or T47D cells (Fig. 5A), and CDCP1 knockdown decreased Met protein expression (Fig. 5B). These findings suggest that CDCP1 functionally interacts with Met, even in cancer cells. Boyden chamber assays revealed that CDCP1 knockdown significantly inhibited HGF-induced cell migration (Fig. 5C) and invasion (Fig. 5D). The inhibition of invasive activity was rescued by re-expression of wild-type CDCP1, but not by the CDCP1-CG or CDCP1-PR mutant. Furthermore, HGF-induced formation of lamellipodia, a membrane structure required for cell migration, was also inhibited by CDCP1 knockdown and rescued only by wild-type CDCP1 (Fig. 5, E and F). These observations demonstrate that CDCP1–Src in lipid rafts plays crucial roles in HGF-induced dynamic cell migration and in the invasion of cancer cells.

**Fig. 5.**
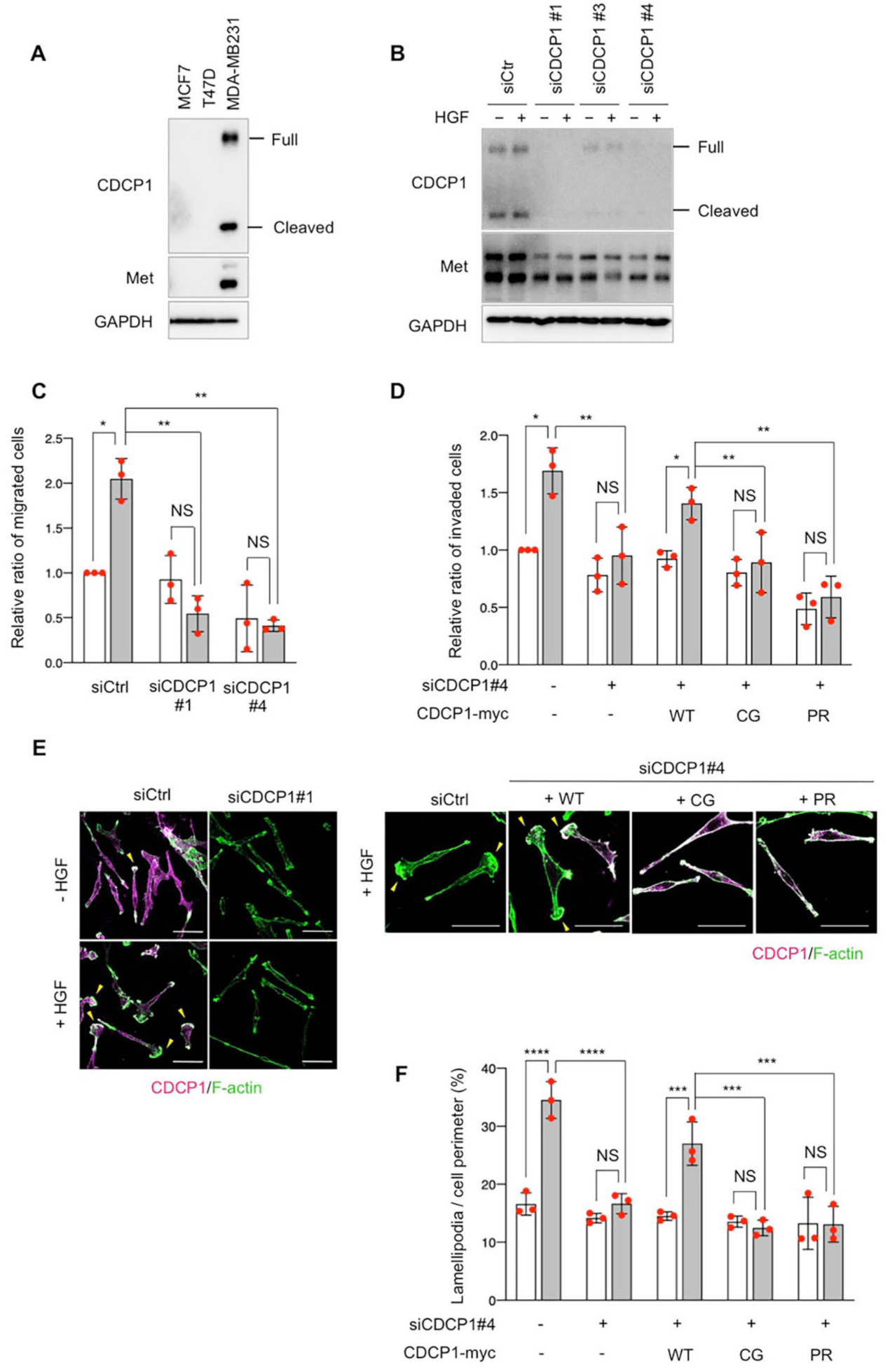
CDCP1 is required for HGF-dependent cancer invasion. (**A**) Total lysates from MCF7, T47D, and MDA-MB231 cells were subjected to immunoblotting using the indicated antibodies. (**B**) MDA-MB231 cells treated with or without HGF (100 ng/ml) were transfected with the indicated CDCP1 siRNAs, and total cell lysates were subjected to immunoblotting using the indicated antibodies. (**C**) The in vitro migration activities of MDA-MB231 cells treated with control and siRNAs were examined by performing transwell assays in the presence or absence of HGF. (**D**) The in vitro invasion activity of control- and siRNA-treated MDA-MB231 cells was examined by performing Matrigel transwell assays in the presence or absence of HGF. Rescue experiments were also performed by re-expressing wild-type CDCP1-myc, CDCP1-CG-myc, and CDCP1-PR-myc. (**E, F**) Formation of lamellipodia in control- and siRNA-treated MDA-MB231 cells was examined by immunofluorescent analysis for F-actin in the presence or absence of HGF (left panels). Yellow arrowheads indicate lamellipodia. Rescue experiments were also performed by re-expressing wild-type CDCP1-myc, CDCP1-CG-myc, and CDCP1-PR-myc (upper right panels). The ratios of the length of lamellipodial membrane to the total cell perimeter were calculated from at least 30 cells of each cell type (lower right graph). The scale bars indicate 50 µm. The mean ratios ± SDs were obtained from three independent experiments. *, *P* < 0.05; **, *P* < 0.001; ***, *P* < 0.001; ****, *P* < 0.0001; NS, not significantly different; two-way ANOVA.

### CDCP1 contributes to induction of compensatory renal growth

Finally, to verify the aforementioned model *in vivo*, we investigated the role of the CDCP1-mediated Met–STAT3 pathway in compensatory renal growth. For this study, we generated Cdcp1-knockout mice using the CRISPR-Cas9 system (Fig. S7, A–C). The Cdcp1-knockout (*Cdcp1*^−/−^) mice grew normally and did not show any overt phenotype under normal conditions (Fig. S7, C–E), as reported previously (Spassov, Wong, Wong, Reiter, & Moasser, 2013). Eight weeks after UNX, the remaining kidney was enlarged in wild-type and heterozygous (*Cdcp1*^+/−^) mice (Fig. 6A). Kidney/body weight ratios in wild-type and *Cdcp1*^+/−^ mice were elevated to approximately 136% and 135% compared to those in sham-operated mice, respectively (Fig. 6B). In contrast, the kidney/body weight ratio elevation was significantly smaller in *Cdcp1*^−/−^ mice (approximately 121%) (Fig. 6, A and B). Because compensatory renal growth is achieved via the expansion of proximal renal tubules (Anderson, 1967; Chen et al., 2015), we visualized the renal tubules using a fluorescein-labeled specific lectin (LTL-FITC). Wild-type mice exhibited prominent thickening of LTL-positive proximal renal tubules (to approximately 148%), whereas this thickening was suppressed in *Cdcp1*^−/−^ mice (approximately 117%) (Fig. 6, C and D). These data confirm that compensatory growth of proximal renal tubules was suppressed in *Cdcp1*^−/−^ mice.

**Fig. 6.**
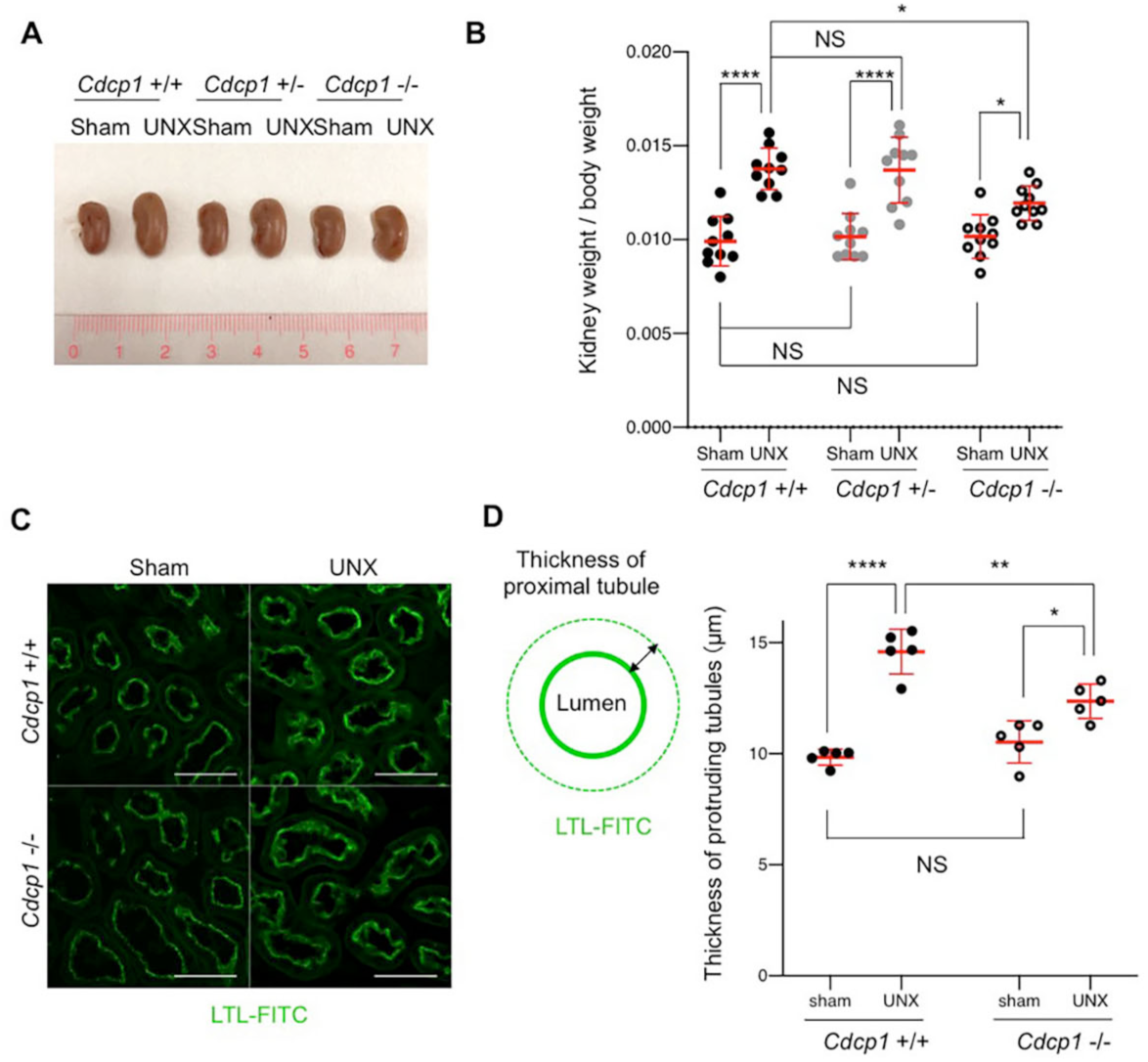
Compensatory renal growth after UNX is suppressed in *Cdcp1*-knockout mice. (**A**) *Cdcp1* wild-type (+/+), heterozygous (+/–), and homozygous knockout (–/–) mice at 8 weeks of age were subjected to left UNX or a sham operation. Regenerative growth of the remaining left kidney was analyzed 8 weeks after operation and increases in remaining kidney/body weight ratios were assessed. (**B**) The mean ratios ± SDs were obtained from 10 mice per group. (**C**) The remaining kidney was removed from UNX- or sham-operated mice, and proximal tubules were stained with FITC-LTL (green). The scale bars indicate 50 µm. (**D**) The proximal tubule thickness was determined by the length of FITC-LTL-stained epithelial cell layer (n = 100). The mean ratios ± SDs were obtained from 5 mice per group. *, *P* < 0.05; **, *P* < 0.01; ****, *P* < 0.0001; NS, not significantly different; two-way ANOVA.

### CDCP1 induces compensatory renal growth by activating Met–STAT3 signaling

To address the cause of defective renal growth in *Cdcp1*^−/−^ mice, we analyzed the remaining kidneys at earlier stages of compensatory growth (within 4 days after UNX) because the expression of HGF and Met is transiently upregulated in renal tissues within 12 h (Ishibashi et al., 1992; Nagaike et al., 1991). The mass of the remaining kidney immediately increased in wild-type mice (Fig. 7A), whereas acute enlargement of the kidney tended to be delayed in *Cdcp1*^−/−^ mice. Immunofluorescence analysis revealed that activation of Met (pY1234/1235) and STAT3 (pY705) occurred in wild-type renal tubules 12 h after UNX, while the activation of both signals was appreciably attenuated in *Cdcp1*^−/−^ renal tubules (Fig. 7, B and C). Activation of CDCP1 (pY734) also occurred in renal tubules in a manner similar to that of Met and STAT3 (Fig. S8A). Notably, activated Met, STAT3, and CDCP1–Src were co-localized in a subset of intracellular small vesicles that were positive for EEA1 (Fig. S8, B–D), a marker of early endosomes, supporting the functional interactions among these signaling molecules. These results suggest that the CDCP1–Src axis mediates HGF–Met–STAT3 signaling even in proximal renal tubules.

**Fig. 7.**
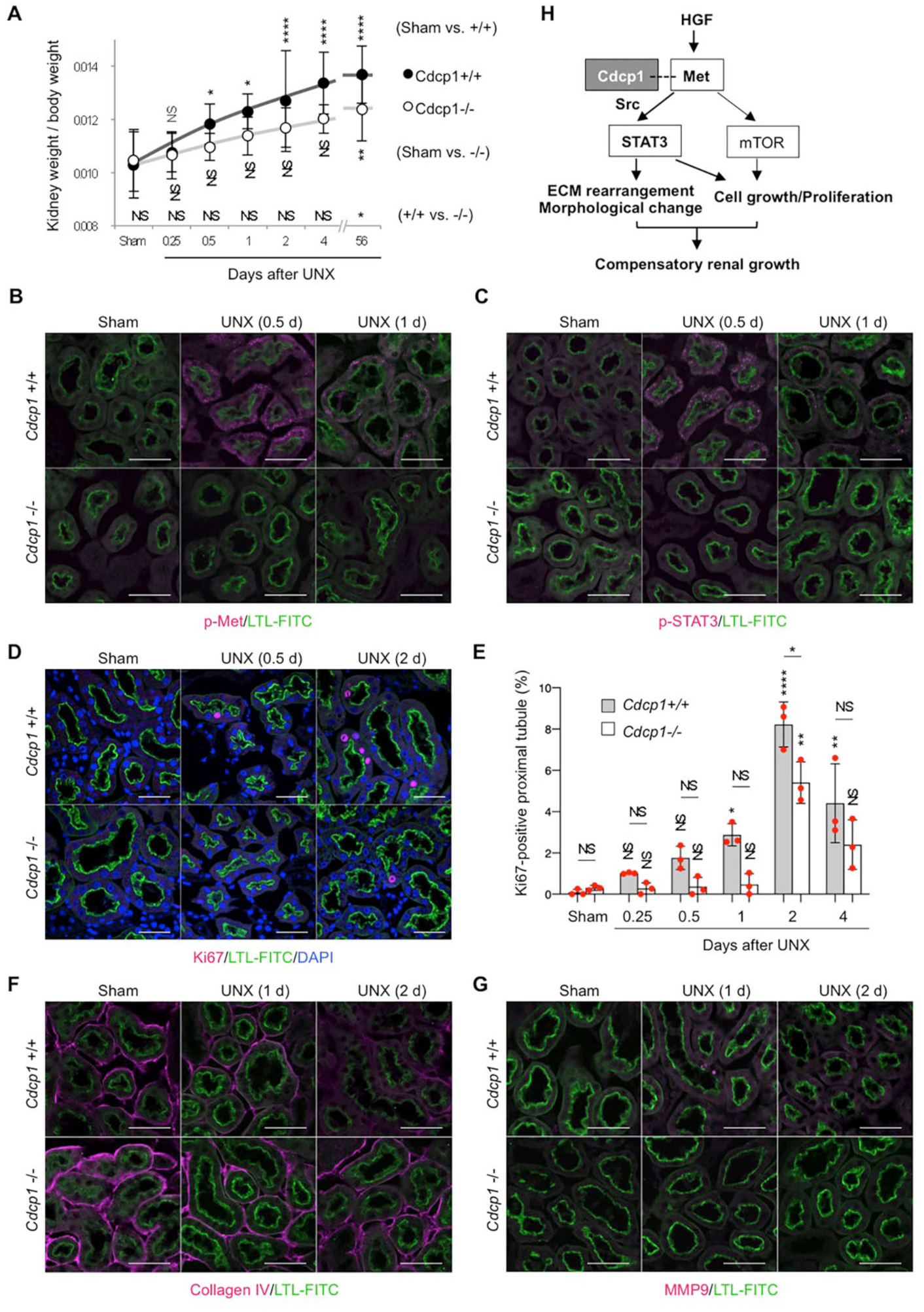
Met–STAT3 signaling is attenuated in *Cdcp1*-knockout mice. (**A**) *Cdcp1* wild-type (+/+) and homozygous knockout (–/–) mice at 8 weeks of age were subjected to left UNX or a sham operation. Compensatory renal growth of the remaining left kidney was analyzed at the indicated time points after operation and increases in remaining kidney/body weight ratios were assessed. The mean ratios ± SDs were obtained from at least 6 mice per group. (**B–D, F, G**) The remaining kidney after UNX was subjected to microscopic immunofluorescence analysis with specific antibodies against Met pY1234-1235 (**B**), STAT3 pY705 (**C**), Ki67 (**D**), collagen IV (**F**), MMP9 (**G**), and Alexa Fluor 594-conjugated secondary antibody (magenta). Proximal tubules were visualized by staining with FITC-LTL (green). The scale bars indicate 50 µm. Ki67-positive proximal tubules (%) were estimated by calculating the ratio of Ki67-stained tubules to the total number of tubules (n > 100) (**E**). The mean ratios ± SD were obtained from 3 mice per group. *, *P* < 0.05; **, *P* < 0.01; ****, *P* < 0.0001; NS, not significantly different; two-way ANOVA was calculated compared to sham operated control. (**H**) Schematic model of HGF-induced adaptive renal regeneration. CDCP1–Src regulated Met–STAT3 signaling leading to compensatory renal growth through the induction of ECM rearrangement and cell growth/proliferation.

We also analyzed cell proliferation by immunostaining for Ki67. The ratios of Ki67-positive cells in wild-type proximal tubules gradually increased and peaked 2 days after UNX (Fig. 7, D and E). However, in *Cdcp1*^−/−^ mice, the appearance of Ki67-positive cells was delayed, and the overall number of proliferating cells decreased. These results suggest that HGF-induced cell proliferation through Met-STAT3 signaling is also attenuated in *Cdcp1*^−/−^ mice, thereby retarding the onset of compensatory renal growth.

We further examined the effects of CDCP1 loss on the surrounding environment of proximal tubules after UNX. Before UNX, collagen IV was observed around the proximal tubules in both wild-type and *Cdcp1*^−/−^ mice (Fig. 7F). In wild-type mice, collagen IV gradually degraded 24 h after UNX, but this degradation was attenuated in *Cdcp1*^−/−^ mice. Consistently, MMP2 and MMP9 were upregulated and occasionally concentrated in vesicle-like structures in proximal tubules 24 h after UNX (Fig. 7G and Fig. S8E). However, these phenomena were barely detectable in *Cdcp1*^−/−^ proximal tubules. These observations suggest that secretory MMPs remodel extracellular matrix for progression of compensatory renal growth downstream of CDCP1–Src-mediated Met-STAT3 signaling.

## Discussion

To address the regulatory mechanisms underlying the multifaceted functions of HGF, i.e., induction of invasive growth phenotypes, we dissected the mechanism of HGF signaling using MDCK cysts and identified Src and its scaffold CDCP1 as critical elements of HGF–Met signaling. Upon stimulation with HGF, CDCP1 concentrated temporally at the tip of protruding cells. Considering that CDCP1 expression is induced by HGF (Gusenbauer, Vlaicu, & Ullrich, 2013) or by epidermal growth factor (EGF)-mediated activation of the MAPK pathway (Adams et al., 2015; Dong et al., 2012), the observed upregulation of CDCP1 might be caused by activation of the HGF–Met–MAPK pathway. We also showed that continuous CDCP1 upregulation induced the formation of multiple cell protrusions and cell growth/proliferation. However, when CDCP1 expression was halted by removing Dox from the media, cells in the protruding cord-like structures recovered epithelial features, resulting in the formation of a luminal structure (Fig. S9A). This finding suggests that the temporal CDCP1 upregulation by growth factors such as HGF might trigger the initial phase of morphogenesis, specifically cell protrusion extension and growth, and that the coordinated negative feedback regulation of CDCP1 might be involved in establishing the compound tubular/acinar epithelial system.

In this study, we demonstrated that activation of the CDCP1–Src complex in lipid rafts was required to induce HGF responses in MDCK cysts. Src activation by CDCP1 in lipid rafts induced STAT3 phosphorylation at Tyr705 (Bromberg et al., 1998; C. L. Yu et al., 1995), leading to the upregulation of MMPs and mitogenic factors such as Myc and Cyclin D1, which promote cell protrusion extension and cell proliferation, respectively. These findings suggest that STAT3 activation via CDCP1-Src in lipid rafts is crucial for inducing invasive growth phenotype. We also found that a physical association between cleaved/activated CDCP1 and Met was required for STAT3 activation. Because STAT3 interacts directly with activated Met through an SH2 domain (Boccaccio et al., 1998), it is likely that the cleaved form of CDCP1 binds Src and interacts with the Met–STAT3 complex in lipid rafts to enable efficient STAT3 phosphorylation by Src (Fig. 4G). Hence, this focal integration of the CDCP1–Src axis with the Met–STAT3 complex might be a key event in controlling HGF responses. However, CDCP1 can also interact with other transmembrane receptors, including HER2, EGFR, integrin β1, and E-cadherin (Alajati et al., 2015; Casar et al., 2014; Law et al., 2013; Law et al., 2016). For example, HER2 associates stably with CDCP1 and forms a heterodimer complex on the plasma membrane; this interaction enhances Src activity in an EGF-independent manner (Alajati et al., 2015). Together with our results, these findings suggest that CDCP1 can also serve as a general regulator of membrane receptor signaling, which involves Src activation.

CDCP1 upregulation has been implicated in tumor progression (He et al., 2016; Ikeda et al., 2009; Miyazawa et al., 2010), particularly cancer invasion and metastasis (He et al., 2016; Liu et al., 2011; Uekita & Sakai, 2011; Wright et al., 2016). In this study, we showed that CDCP1 and Met were co-upregulated in highly invasive breast cancer MDA-MB231 cells, but not in non-invasive MCF7 or T47D cells, suggesting that a functional interaction occurred between them. Indeed, CDCP1 knockdown downregulated Met and inhibited HGF-dependent invasive activity. Because Src binding sites in CDCP1 and lipid rafts were requited for the promotion of HGF-dependent invasive activity, CDCP1-mediated Src activation in lipid rafts likely contributes to cancer invasion in a manner similar to that in MDCK cysts. Interestingly, we also found that CDCP1 upregulation in MDCK cysts induced the expression of cytokeratin 14 (Fig. S9B), a marker of collective cancer invasion (Cheung, Gabrielson, Werb, & Ewald, 2013). This finding suggests a potential role for the CDCP1–Src axis, even in collective cancer invasion. In support of these observations, The Cancer Genome Atlas (TCGA) database analysis revealed a significant correlation between CDCP1^high^/MET^high^ groups and poor prognoses in patients with breast or kidney cancer (Fig. S10, A and B). Therefore, further analyses of the functions of the CDCP1–Src–Met–STAT3 pathway in a wide range of cancers could reveal new therapeutic targets for treating some malignant cancers.

HGF is immediately upregulated after the loss of kidney mass and plays important roles in compensatory growth by inducing cell proliferation and anti-apoptotic effects (Matsumoto & Nakamura, 2001). In this study, we uncovered the contribution of CDCP1–Src to HGF-induced compensatory renal growth using Cdcp1-deficient mice. We showed that activation of Met and STAT3 in the renal tubule cells following UNX was significantly attenuated in Cdcp1-deficient mice. Furthermore, MMP2/9 upregulation and basement membrane collagen IV degradation (Tan & Liu, 2012) were detected in the renal tubule cells after UNX (Fig. S8F). These observations are consistent with those of our *in vitro* study in MDCK cysts, supporting the crucial role of the functional integration of CDCP1–Src with Met–STAT3 pathways during compensatory renal growth. Given that CDCP1 expression is temporally upregulated under pathological conditions such as tissue injury, hypoxia, and cancer recurrence (He et al., 2016), it is likely that upregulated CDCP1 might acutely amplify the HGF signaling required for regenerative organ growth in critical situations. Furthermore, because STAT3 is an important regulator of the regeneration of other organs such as the liver, muscles, and intestines (Lindemans et al., 2015; Taub, 2004; Tierney et al., 2014), it is possible that the temporal upregulation of CDCP1 widely contributes to the promotion of organ regeneration by inducing focal activation of the Src–STAT3 pathway.

In conclusion, we showed that CDCP1 served as a critical regulator of HGF-induced invasive growth phenotypes by focally integrating Src and the Met–STAT3 pathway (Fig. 7H). Our discovery of the CDCP1–Src axis as a new component of HGF–Met signaling provides insights into the regulatory mechanisms underlying the multifaceted functions of HGF during regenerative growth and cancer invasion, and might contribute to the development of more promising therapies for malignant cancers that are associated with the co-upregulation of CDCP1 and Met.

## Materials and Methods

### Cell culture

MDCK (type-I), HEK293T, MDA-MB231, MCF7, and T47D cells were cultured in Dulbecco’s modified Eagle’s medium (DMEM) supplemented with 10% fetal bovine serum (FBS) at 37 °C in a 5% CO_2_ atmosphere. Three-dimensional culture was performed using type-I collagen (Cellmatrix Type I-A, Nitta Gelatin) according to the manufacturer’s protocol. Type I-A collagen (3 mg/ml) was neutralized with reconstitution buffer (2.2% NaHCO_3_, 0.05 N NaOH, and 200 mM HEPES) and diluted with 5× DMEM (Gibco). MDCK cells (1.5 × 10^5^ cells/ml of collagen gel) were combined with collagen in DMEM supplemented with 5% FBS, and then polymerized at 37°C in a 5% CO_2_ atmosphere. The medium was replaced every 2 days. A collagen gel containing 2% type-I DQ-collagen (Molecular Probes) was used for the DQ-collagen degradation assay.

### Mice

*Cdcp1*-knockout mice were generated in a C57BL/6N background using the CRISPR/Cas9 system. Animals were housed in environmentally controlled rooms at the animal experimentation facility of Osaka University. All animal experiments were conducted according to the guidelines of the Osaka University committee for animal and recombinant DNA experiments and were approved by the Osaka University Institutional Review Board. The guide RNA (gRNA) and primer sequencess used for genotyping are shown in Supplementary Table 3 (see also Supplementary Fig. 9a).

### Antibodies and inhibitors

For this study, antibodies against CDCP1 (4115), CDCP1 pY734 (9050), STAT3 (9132), STAT3 pY705 (9145), Myc-tag (2276), Met pY1234/1235 (3077), and Met (8198) were purchased from Cell Signaling Technologies. Anti-CDCP1 antibody (LC-C172540) was from LSBio. Antibodies against SFK (sc-18, clone SRC2), ERα (sc-542, clone MC-20), GAPDH (sc-32233, clone 6C5), and MMP2 (sc-10736) were from Santa Cruz Biotechnology. Anti-Src pY418 (44-655G), anti-Src pY529 (44-662G), and anti-Ki67 (14-5698-82) antibodies were from Thermo Fisher Scientific. Anti-Src (OP07, clone Ab-1), anti-phosphotyrosine (05-1050, clone 4G10), and anti-MMP9 (444236) antibodies were from Millipore. Anti-collagen IV antibody (ab6586) was purchased from Abcam. Anti-laminin antibody (L9393) was from Sigma. Anti-keratin 14 antibody (PRB-155P, clone AF64) was from Covance. Several inhibitors were used in this study. S3i-201 (573102), Stattic (573099), JAK inhibitor 1 (420099), c-Met/Ron dual kinase inhibitor (448104), and Rac1 inhibitor (553502) were purchased from Calbiochem. Marimastat (M2699) was from Sigma. Y27632 (257-00511) was from Wako Chemicals. Other inhibitors listed in Supplementary Table 1 were obtained from the Screening Committee of Anticancer Drugs.

### Immunoblotting and immunoprecipitation

For two-dimensional culture, cells were lysed in n-octyl-β-D-glucoside (ODG) buffer [20 mM Tris-HCl (pH 7.4), 150 mM NaCl, 1 mM EDTA, 1 mM Na_3_VO_4_, 20 mM NaF, 1% Nonidet P-40, 5% glycerol, 2% ODG and a protease inhibitor cocktail (Nacalai Tesque)], and immunoblotting was performed. For three-dimensional culture, the cyst-containing collagen matrix was incubated with HBS buffer (10 mM HEPES [pH 7.3], 140 mM NaCl, 4 mM KCl, 1.8 mM CaCl_2_ and 1 mM MgCl_2_) containing 0.1% collagenase (Roche) at 37 °C. Cysts were harvested by centrifugation and lysed in SDS sample buffer (50 mM Tris-HCl [pH 6.8], 2% SDS, 100 mM NaCl, 1 mM EDTA, 1 mM Na_3_VO_4_, 20 mM NaF and 5% sucrose) before immunoblotting. For immunoprecipitation assays, the cells were lysed in ODG buffer and the lysates were incubated with an anti-Myc-tag antibody (2276, Cell Signaling Technology). Immunoprecipitated proteins were pulled down with protein A-sepharose (GE Healthcare) for immunoblotting. Horseradish peroxidase-conjugated anti-mouse or anti-rabbit IgG (Zymed) was used as the secondary antibody. All immunoblots were visualised and quantitated using a Luminograph II System (Atto). Silver staining was performed using the Silver Stain MS Kit (Wako Chemicals).

### DRM fractionation

Cells were lysed in homogenization buffer (50 mM Tris-HCl [pH 7.4], 150 mM NaCl, 1 mM EDTA, 1 mM Na_3_VO_4_, 20 mM NaF, 0.25% Triton X-100, and protease inhibitor cocktail) and separated on a discontinuous sucrose gradient (5–35–40%) by ultracentrifugation at 150,000 × *g* for 12 h at 4°C using an Optima L-100XP centrifuge equipped with a SW55Ti rotor (Beckman Coulter). Eleven fractions were collected from the top of the sucrose gradient.

### Microarray and quantitative real-time PCR

For microarray analysis, total RNA was isolated from MDCK cysts using the Sepasol-RNA Kit (Nacalai Tesque). Microarray analysis was performed on a G2505C Microarray Scanner (Agilent Technologies) using the Canis (V2) Gene Expression Microarray 4×44K (Agilent Technologies). Microarray data were subjected to gene ontology (GO) analysis and upstream regulator analysis using the IPA Program (Qiagen, http://www.qiagenbioinfomatics.com/). For quantitative real-time PCR analysis, cDNA was prepared from RNA using the Transcriptor First Strand cDNA Synthesis Kit (Roche) according to the manufacturer’s instructions. Real-time PCR was performed on a 7900HT Fast Real-Time PCR System (Applied Biosystems) using the Thunderbird qPCR Mix (Toyobo). Total RNA expression was normalized to expression of the reference gene, GAPDH. The sequences of the primers used for this analysis are listed in Table S2.

### Immunofluoresce microscopy

For two-dimensional culture, cells were grown on coverslips coated with type-I collagen, fixed with 4% paraformaldehyde (PFA) and permeabilized with phosphate-buffered saline (PBS) containing 0.03% Triton X-100. For three-dimensional culture, cysts embedded within the collagen matrix were fixed with 4% PFA and permeabilized with PBS containing 0.5% Triton X-100. Permeabilized cells and cysts were blocked with 1% bovine serum albumin (BSA) and incubated with primary antibodies, and then incubated with Alexa Fluor 488/594-phalloidin (Molecular Probes). For the kidney sections, kidneys were prepared by perfusion fixation with 4% PFA and dissected. The fixed kidneys were embedded in OCT compound (Sakura Finetek), sectioned, and mounted on glass slides. The kidney sections were blocked with Blocking One (Nacalai Tesque), incubated with primary antibodies, and then incubated with Alexa Fluor 488/594-conjugated secondary antibodies (Molecular Probes) and the fluorescein-conjugated proximal tubule marker, FITC-LTL (FL-1321, Vector Laboratories). Nuclei were counterstained with 4’,6-diamidino-2-phenylindole (DAPI). Immunostained objects were observed under an FV1000 confocal microscope (Olympus). For time-lapse observations of three-dimensional cultures, cysts embedded within in collagen matrix were observed under a FV1200 confocal microscope (Olympus).

### CRISPR/Cas9-based generation of CDCP1-knockout MDCK cells

MDCK cells were co-transfected with the pSilencer1.0-U6 plasmid containing target-gRNA, and the pMJ920 plasmid (Addgene) that co-expresses Cas9 and EGFP using the MDCK Cell Avalanche Transfection Reagent (EZ Bioscience) per the manufacturer’s protocol. Three days after transfection, single EGFP-positive cells were isolated with a FACSAria III flow cytometer (BD Biosciences). Knockout of the *Canis CDCP1* gene was confirmed by immunoblotting. The gRNA and primer sequences used for genotyping are shown in Table S3 (see also Fig. S2B).

### Plasmid construction and gene transfer

CDCP1, CDCP1 deletion mutants, STAT3, and Met were generated by PCR using human cDNA as the template and subcloned into the pCX4 retroviral plasmid (generously donated by Dr. Akagi) (Akagi, Sasai, & Hanafusa, 2003). CDCP1 mutants (K365A–R368A–K369A, C689G–C690G and Y734F) and STAT3-Y705F were generated by mutagenesis PCR using KOD-Plus polymerase (Toyobo). CDCP1 and its respective mutants were subcloned into either the pEGFP-N1 or pmCherry-N1 plasmid (Clontech) and then subcloned into the pRetroX-TRE3G retroviral plasmid (Clontech). Src–MER and STAT3–MER were constructed by first amplifying part of the gene encoding estrogen receptor (MER, amino acids 281–599) and subcloning the amplicon into the pCX4 plasmid. mCherry-CAAX was constructed subcloning the C-terminal region of human KRAS (amino acids 166-189) into the pCX4 plasmid. mCherry-GPI was also subcloned into the pCX4 plasmid (generously donated by Dr. Kiyokawa) (Yagi, Matsuda, & Kiyokawa, 2012). All constructs were confirmed by sequencing. Gene transfer of pCX4 and pRrtroX-TRE3G was carried out by retroviral infection. Retroviral production and infection were performed as described previously (Kajiwara et al., 2014). Small-interfering RNAs (siRNAs) against CDCP1 mRNA (SASI_Hs01_00047185-00047188) were purchased from Merck, and transfected with Lipofectamine RNAiMAX (Thermo Fisher Scientific).

### Migration and invasion assay

BioCoat cell culture permeable supports and Matrigel Invasion Chambers (Corning) were used for the migration and invasion assays, respectively. Cells (0.5 × 10^5^ for migration assays and 1 × 10^5^ for invasion assays) were seeded on inserts and transferred to chambers containing culture media and 0.1% FBS, with or without 100 ng/ml HGF. After incubation at 37°C for 24 h, migrated or invaded cells were fixed with 4% PFA and then stained with 1% crystal violet. Invaded cells were counted on micrographs; in each experiment, cells were counted on five randomly chosen fields. The migration and invasion assays were repeated at least three times.

### Surgical procedures of mice

Compensatory renal growth was induced by right UNX in male mice (8 weeks of age) under anesthesia as previously described (Chen et al., 2015). Renal growth was evaluated by measuring the weight of remaining kidney and the body. Left kidneys of sham-nephrectomized mice were used as controls for UNX mice.

### Kaplan-Meier survival analysis

Clinical and RNA-Seq data from the publicly available TCGA dataset from patients with breast cancers and kidney clear cell carcinoma were used for the analysis of CDCP1 and Met expression in tumor samples. Survival curves were estimated using the Kaplan–Meier method, and the obtained survival curves were compared by using the log-rank test.

### Statistics and reproducibility

For data analyses, unpaired two-tailed *t*-tests were performed to determine the *P*-values. For multiple group comparisons, two-way analysis of variance (ANOVA) was used. A *P*-value of <0.05 was considered to reflect a statistically significant difference. All data and statistics were derived from at least three independent experiments.

## Data availability

Supporting microarray data have been deposited in Gene Expression Omnibus under accession code GSE99375. All other supporting data are available from the corresponding author upon request.

## Acknowledgments

We thank Dr. Yamano and Dr. Takata for consultation on the experiments, Dr. Kiyokawa for providing the mCherry-GPI plasmid, Dr. Saito for mass spectrometry analysis, and the members of Spectrography and Bioimaging Facility (NIBB Core Research Facilities). The inhibitor kit was provided by the Screening Committee of Anticancer Drugs, which is supported by Grant-in-Aid for Scientific Research on Innovative Areas, Scientific Support Programs for Cancer Research, from The Ministry of Education, Culture, Sports, Science and Technology of Japan. This work was supported by a Grant-in-Aid for Young Scientists (26830071, to K.K.), a Grant-in-Aid for Scientific Research (C) (19K07639, to K.K.), a Grant-in-Aid for Scientific Research (B) (19H03504, to M.O.), a Grant-in-Aid for Scientific Research on Innovative Area (19H04962, to M.O.), and a Grant-in-Aid for Scientific Research on Innovative Area (16H01447, to K.A.) from the Ministry of Education, Culture, Sports and Technology of Japan. This work was also supported by the Takeda Science Foundation, and the Extramural Collaborative Research Grant of Cancer Research Institute, Kanazawa University (to K.K).

## Competing interests

The authors declare that they have no conflict of interest.

## Supplementary Materials

### Supplementary Figures

**Fig. S1.**
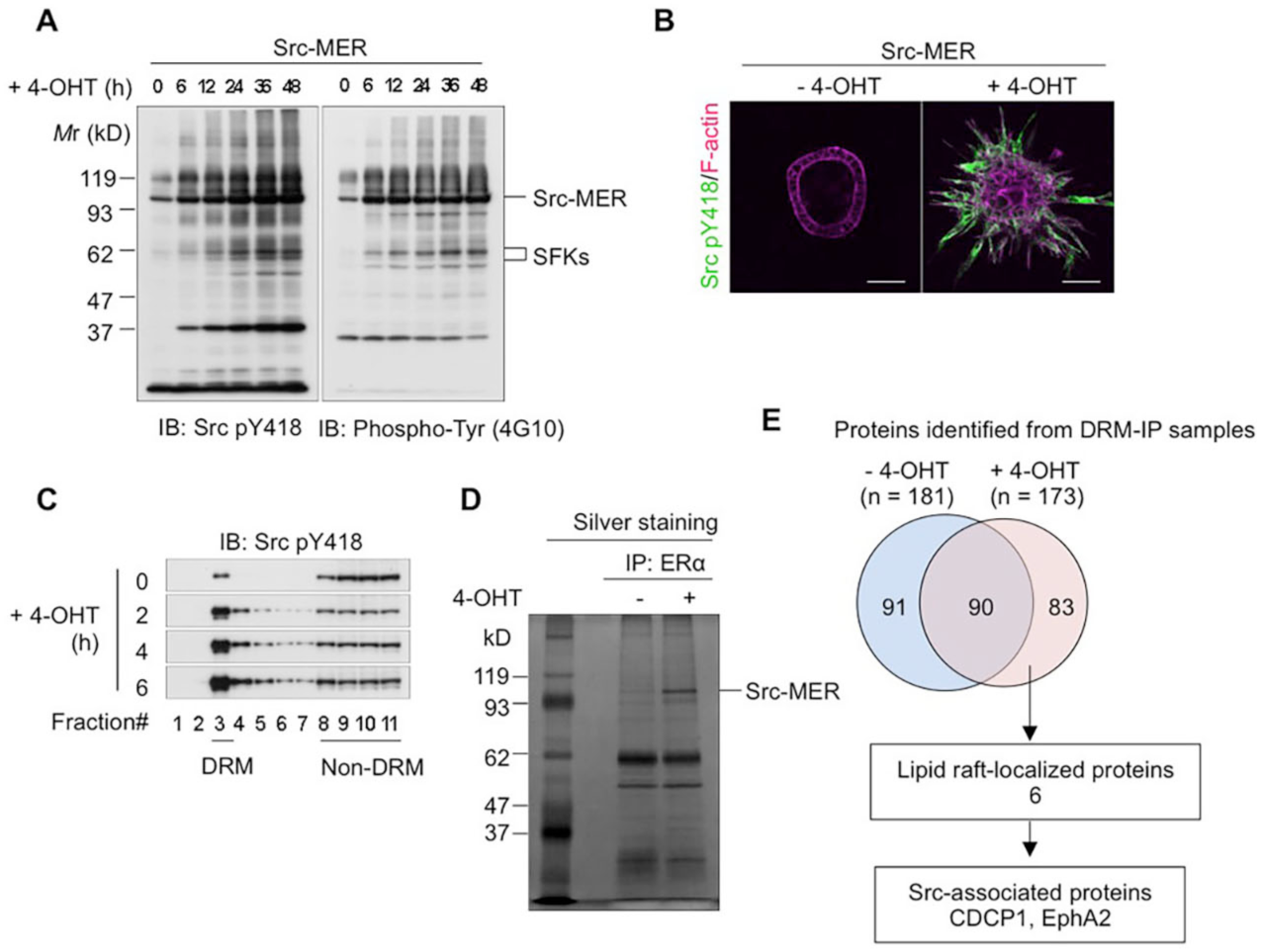
Src associates with CDCP1 in lipid raft fractions. (**A**) Src–MER-overexpressing MDCK cells were incubated with 10 µM 4-OHT for the indicated time periods. Cell lysates were subjected to immunoblotting using the indicated antibodies. (**B**) Src–MER-overexpressing cysts embedded within the collagen matrix were incubated with 10 µM 4-OHT for 2 days. Activated Src was visualized with an anti-Src pY418 antibody (green). Actin filaments were stained with Alexa Fluor 594-phalloidin (magenta). The scale bars indicate 50 µm. (**C**) Src–MER-overexpressing MDCK cells were incubated with 10 µM 4-OHT for the indicated time periods. DRM and non-DRM fractions were separated in a sucrose-density gradient. Aliquots of the fractions were analyzed by immunoblotting with an anti-Src pY418 antibody. **(D)** DRM fractions from Src–MER-activated cells were subjected to immunoprecipitation with an anti-estrogen receptor (ERα) antibody. Immunoprecipitates were confirmed by silver staining. **(E)** Schematic flow diagram of process used to identify Src-associated scaffolding proteins. All proteins in a polyacrylamide gel were trypsinized and subjected to mass spectrometry. Proteins were identified using the SwissProt database, and the number of identified proteins is depicted.

**Fig. S2.**
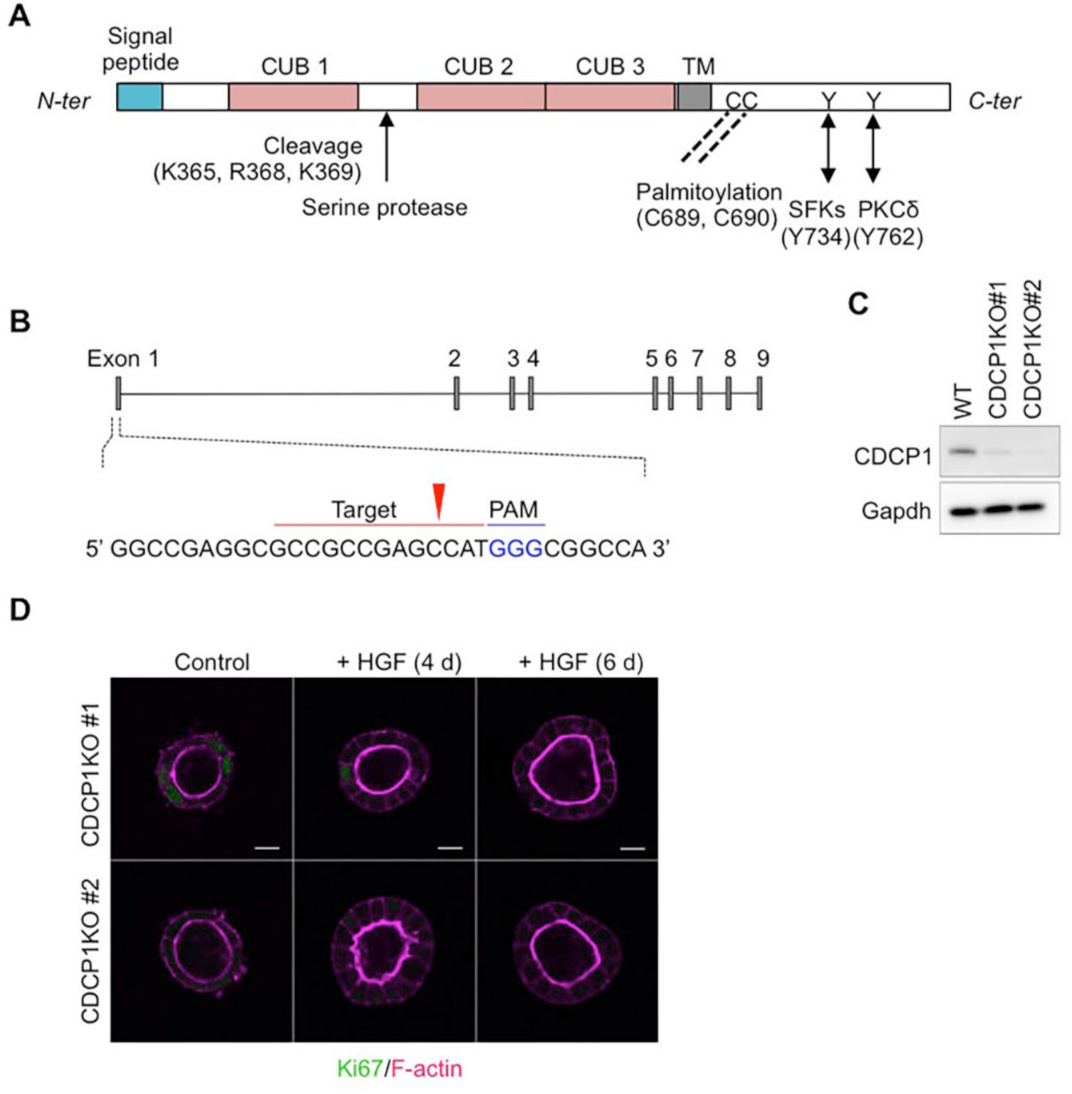
Generation of CDCP1-knockout MDCK cells. (**A**) Schematic illustration of the structure of CDCP1. In the cytoplasmic region, CDCP1 contains a Src-association motif around Tyr734 and two palmitoylation sites (Cys residues 689 and 690), which are required for CDCP1 localization in lipid rafts. Upon Tyr734 phosphorylation by Src, CDCP1 is additionally phosphorylated at Tyr762, resulting in direct association with PKCd, which promotes cell migration. In the extracellular region, CDCP1 contains three CUB domains that are required for protein–protein interactions. CDCP1 also harbors a proteolytic cleavage (shedding) site between the first and second CUB domains and can, thus, be present in either the full-length or cleaved form, depending on the cellular context. TM indicates the transmembrane domain. (**B**) Schematic diagram of CRISPR/Cas9-based generation of CDCP1-knockout MDCK cells. The blue text indicates the PAM sequence and the bold letters indicate the start codon. The red arrowhead indicates the cleavage site. (**C**) Immunoblotting analysis of CDCP1-knockout MDCK cells. Lysates from wild-type and CDCP1-knockout MDCK cells were subjected to immunoblotting using the indicated antibodies. (**D**) CDCP1-knockout MDCK cysts were incubated in the presence of HGF (50 ng/ml) for 4 or 6 days. Ki67 was visualized with an Alexa Fluor 488-conjugated antibody (green) and actin filaments were stained with Alexa Fluor 594-phalloidin (magenta). The scale bars indicate 10 µm.

**Fig. S3.**
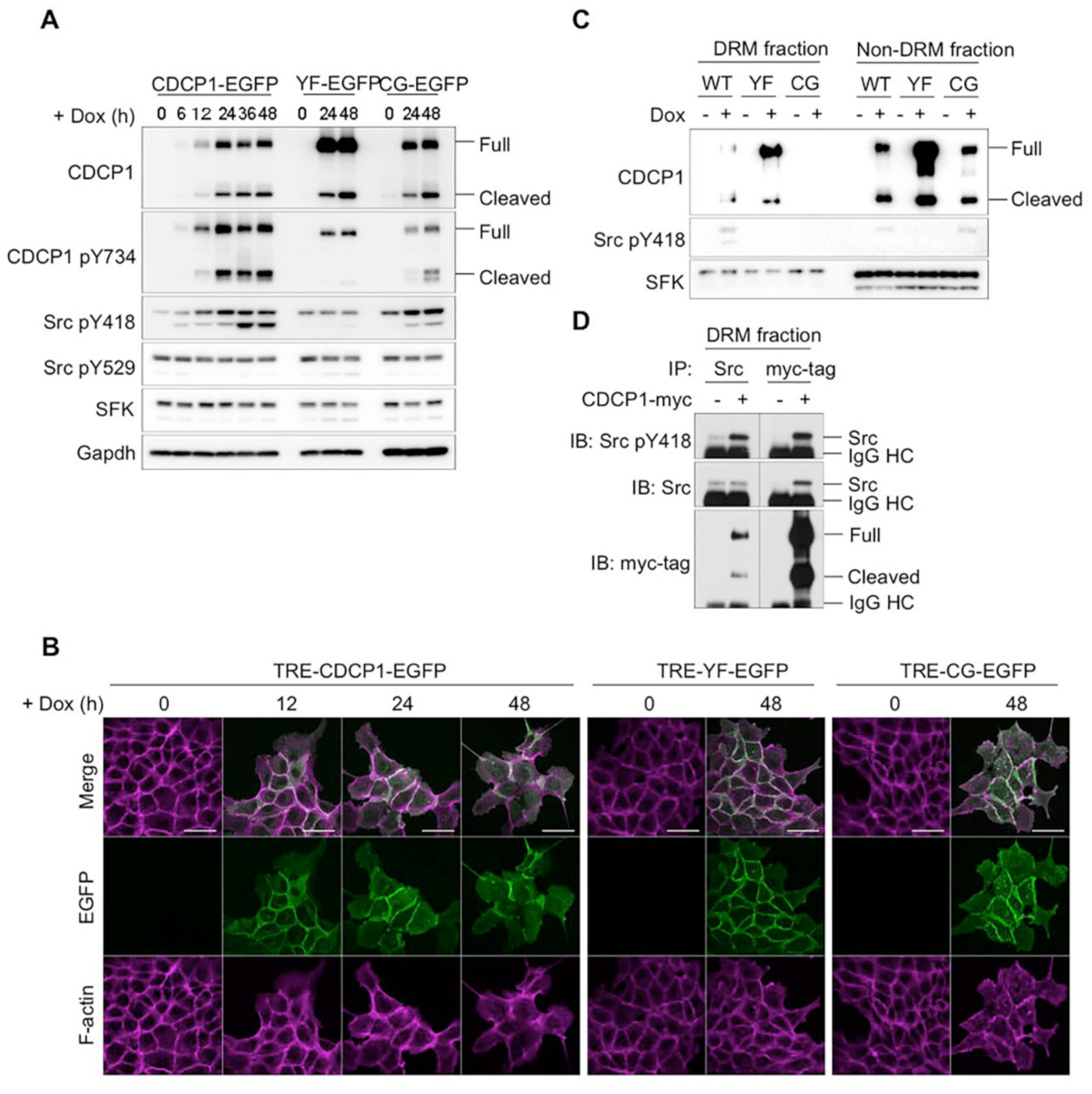
CDCP1 recruits activated Src into lipid rafts. (**A**) TRE–CDCP1–EGFP (CDCP1–EGFP cells)- and mutant (YF-EGFP or CG-EGFP)-harboring MDCK cells were incubated in the presence of Dox (1 µg/ml) for the indicated time periods. The cell lysates were subjected to immunoblotting using the indicated antibodies. (**B**) CDCP1–EGFP cells were incubated in the presence of Dox (1 µg/ml) for the indicated time periods. Actin filaments were stained with Alexa Fluor 594-phalloidin (magenta). The scale bars indicate 50 µm. (**C**) CDCP1–EGFP- and mutant-harboring MDCK cells were incubated in the presence of Dox (1 µg/ml) for 48 h. DRM and non-DRM fractions were separated on a sucrose-density gradient. Aliquots of the fractions were subjected to immunoblotting analysis, using the indicated antibodies. (**D**) DRM fractions of CDCP1–Myc-overexpressing MDCK cells were immunoprecipitated with an anti-Src or anti-Myc tag antibody. Immunoprecipitates were subjected to immunoblotting using the indicated antibodies (HC, heavy chain).

**Fig. S4.**
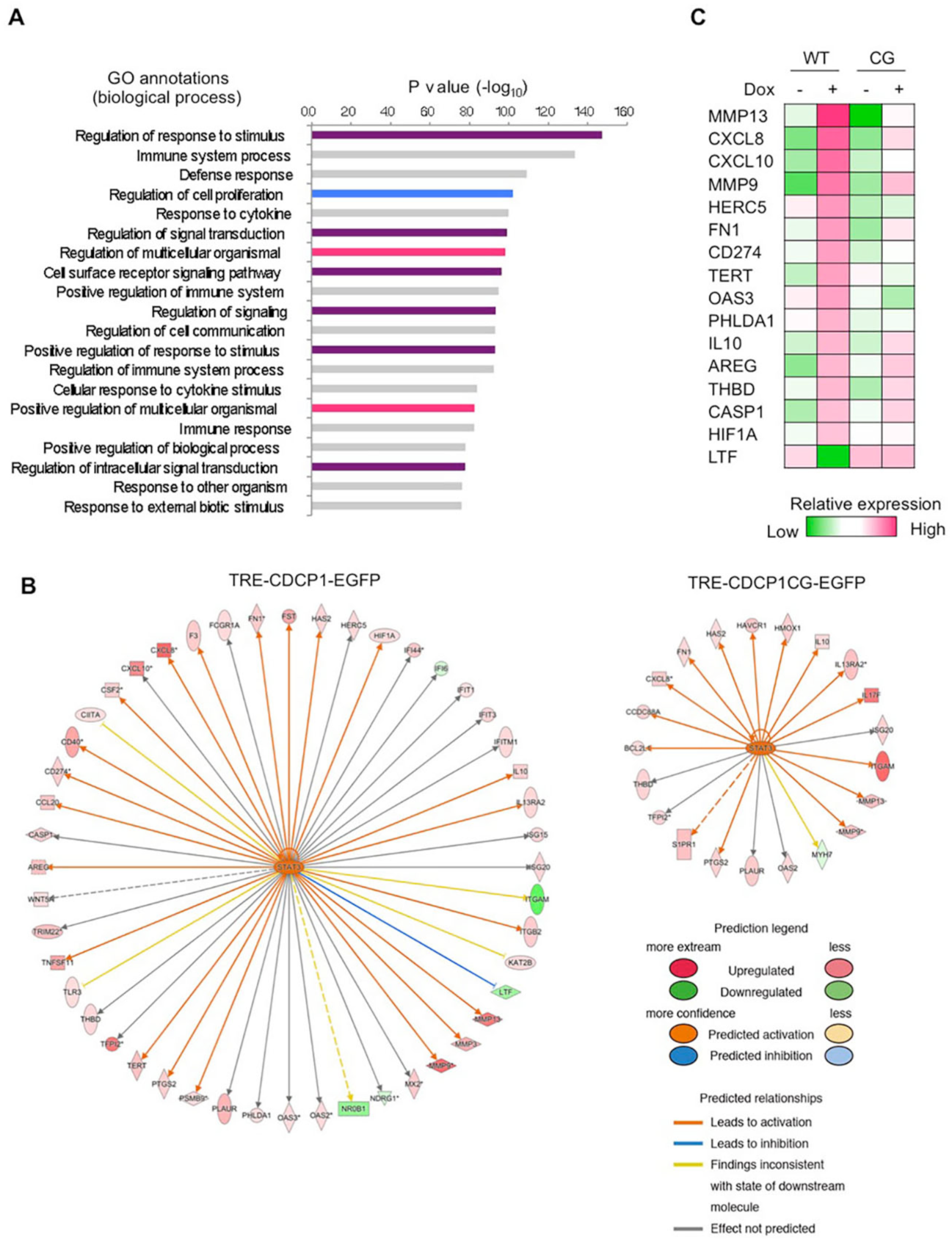
Ingenuity Pathway Analysis of CDCP1-overexpressing cysts. (**A**) GO analysis of CDCP1–EGFP cysts. Magenta bars indicate signaling related to GO annotations. (**B**) Regulatory networks upstream of STAT3 in CDCP1–EGFP and CDCP1–CG-EGFP cysts. Activation Z-scores are presented in Supplementary Table 2. Red and green symbols indicate transcript levels that were upregulated and downregulated by CDCP1 overexpression, respectively. Arrows are colored to indicate activation (orange), inhibition (blue), inconsistency with the state of the downstream molecule (yellow), and unknown effects (grey). (**C**) Heat map representing changes in the expression of STAT3 target genes in CDCP1–EGFP (WT) and CDCP1–CG-EGFP (CG) cysts.

**Fig. S5.**
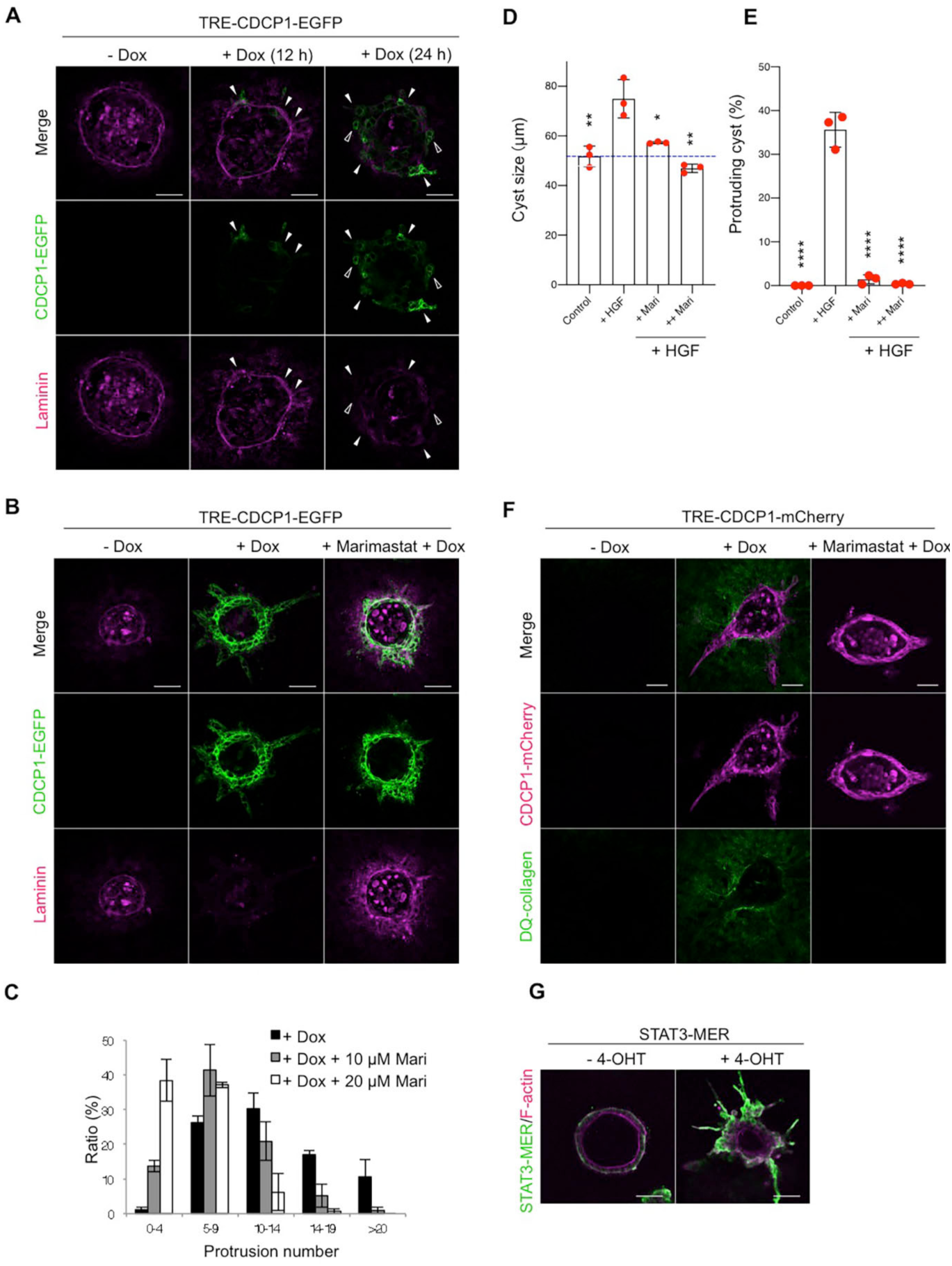
Alterations of extracellular environments by CDCP1 overexpression. (**A**) CDCP1–EGFP cysts embedded within the collagen matrix were incubated with Dox (1 µg/ml) for the indicated time periods. The basement membrane was visualized with an anti-laminin antibody (magenta). The arrowheads indicate protruding cells, and the open arrowheads indicate multi-layered structures. (**B**) CDCP1–EGFP cysts were pretreated with 20 µM marimastat for 2 h and then incubated with Dox (1 µg/ml) for 4 days. The basement membrane was visualized using an anti-laminin antibody (magenta). (**C**) The histogram depicting the percentage of cells with protrusions was calculated by setting the total number of cysts to 100% (n > 150). (**D, E**) MDCK cysts embedded within the collagen matrix were pretreated with marimastat (+ Mari, 10 µM; ++ Mari, 20 µM) for 2 h and then incubated in the presence of HGF (50 ng/ml) for 1 day. (**D**) Diameter (µm) of cysts (n = 100). The dotted blue line indicates the average diameter of non-treated cysts. (**E**) Fraction of the total number of cysts counted (n > 100). The mean ratios ± SDs were obtained from three independent experiments. *, *P* < 0.05; **, *P* < 0.01; **** *P* < 0.0001. Two-way ANOVA was performed relative to the HGF-treated cysts. (**F**) TRE–CDCP1–mCherry-harboring MDCK cells were embedded within a 2% DQ-collagen-containing matrix. CDCP1–mCherry cysts were pretreated with 20 µM marimastat for 2 h and then incubated with Dox (1 µg/ml) for 4 days. DQ fluorescence (green) was visualized under a fluorescent microscope. Scale bars indicate 50 µm. (**G**) STAT3–MER-overexpressing cysts embedded within the collagen matrix were incubated with 1 µM 4-OHT for 4 days. STAT3–MER was visualized with ERα antibody (green) and actin filaments were stained with Alexa Fluor 594-phalloidin (magenta). The scale bars indicate 50 µm.

**Fig. S6.**
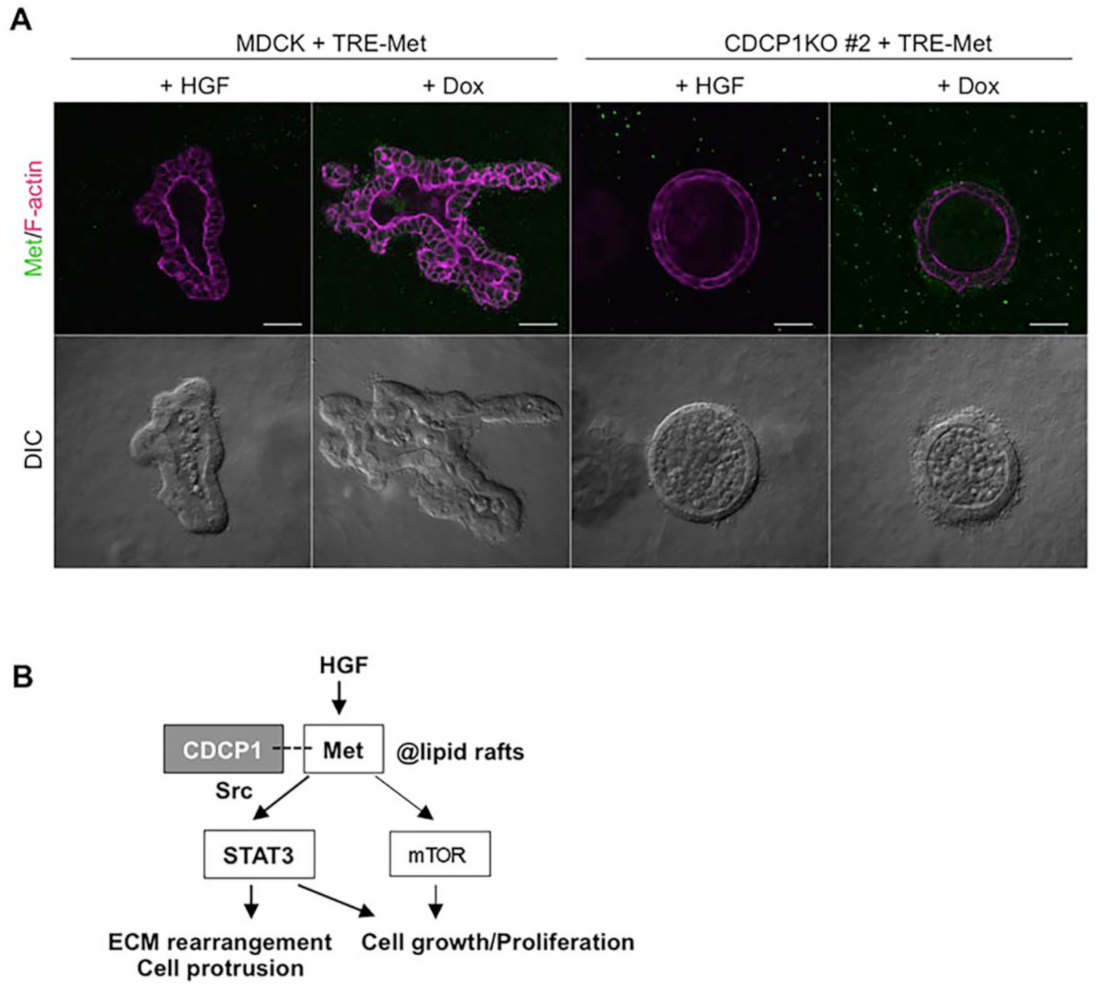
Met-induced morphological changes are inhibited by CDCP1 knockout. (**A**) Wild-type and CDCP1-knockout TRE–Met-harboring MDCK cells embedded within the collagen matrix were incubated with HGF (50 ng/ml) or Dox (1 µg/ml) for 4 days. Met was visualized with an anti-Met antibody (green). Scale bars indicate 50 µm. (**B**) A schematic model of the role of CDCP1–Src in HGF-induced invasive growth. CDCP1–Src activates Met–STAT3 signaling in lipid rafts, leading to invasive growth by inducing ECM rearrangement and cell growth/proliferation. The mTOR pathway also contributes to cell growth.

**Fig. S7.**
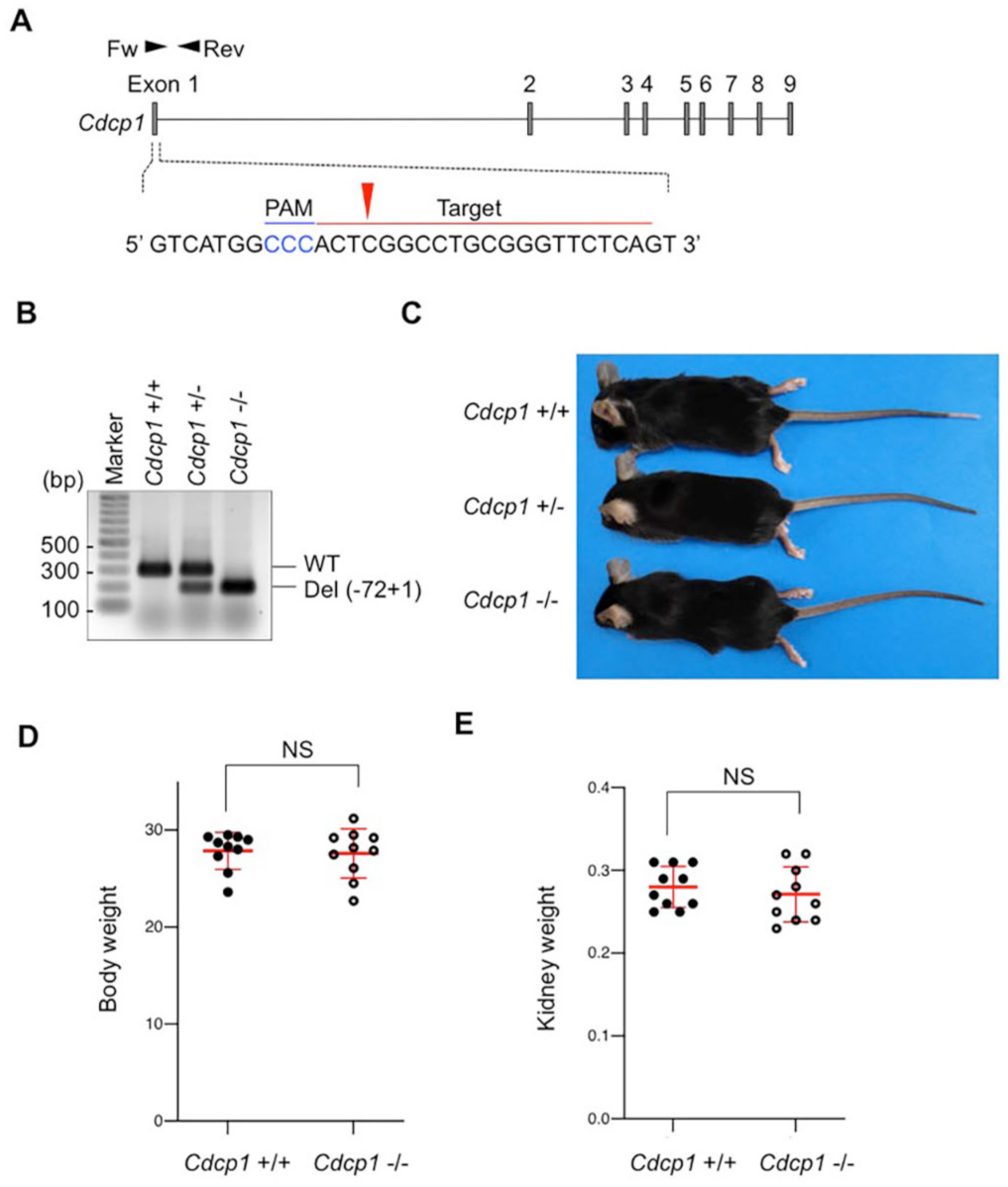
Generation of *Cdcp1*-knockout mice. (**A**) Schematic diagram of CRISPR/Cas9-based generation of *Cdcp1*-knockout mouse. The blue text indicates the PAM sequence and the bold letters indicate the start codon. The red arrowhead indicates the cleavage site. (**B**) PCR verification of deletion of the 1st exon of *Cdcp1* using the forward and reverse primers depicted in panel (**A**). (**C**) Representative picture of wild-type (+/+), *Cdcp1* heterozygous (+/–), and homozygous knockout (–/–) littermate mice at 8 weeks of age. (**D, E**) Whole body (**D**) and left kidney (**E**) weight of wild-type (+/+) and *Cdcp1* homozygous knockout (–/–) mice at 8 weeks of age. The mean ratios ± SDs were obtained from ten mice per group. NS, not significantly different; unpaired two-tailed *t*-test

**Fig. S8.**
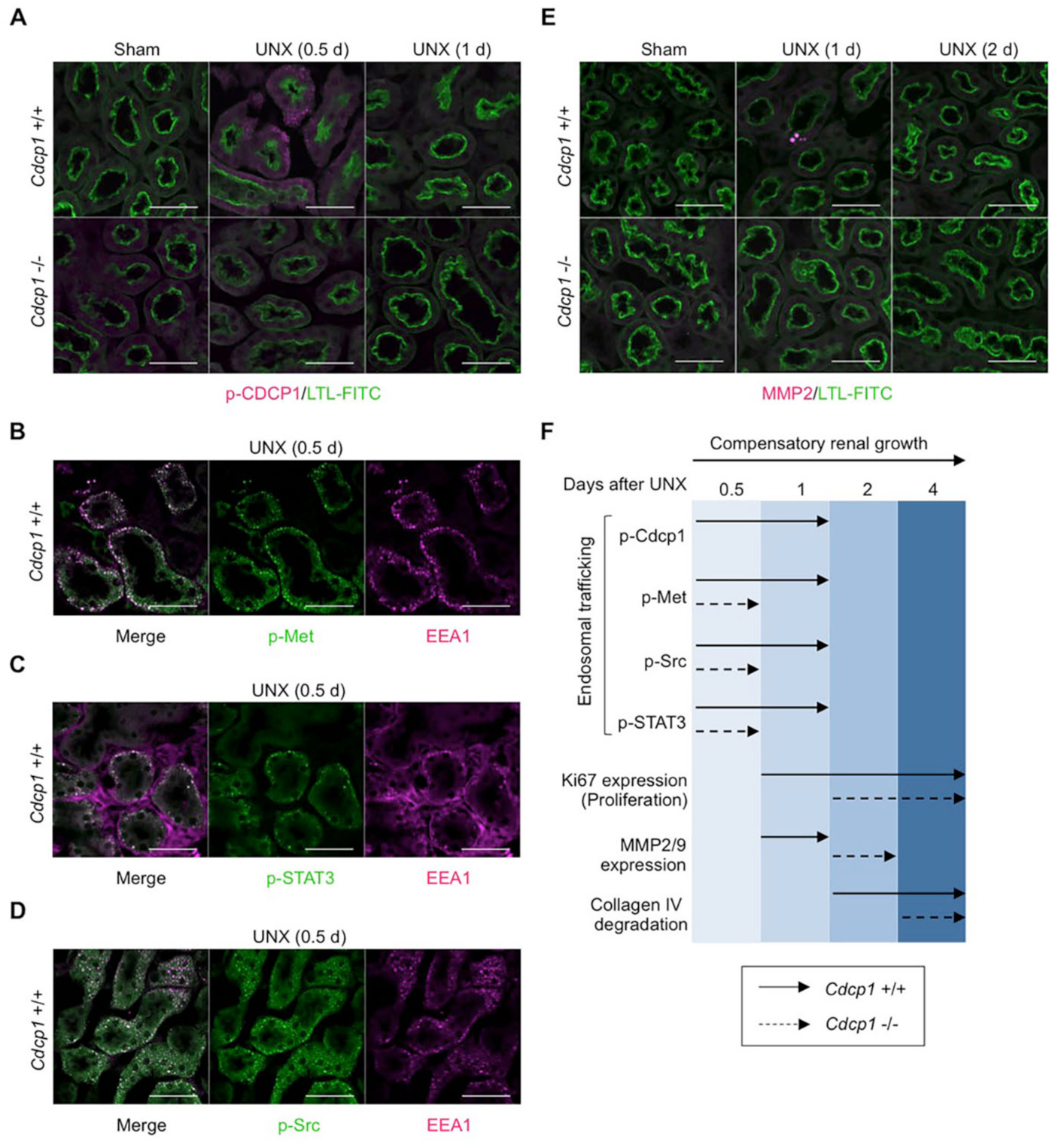
Met–STAT3 signaling is attenuated in *Cdcp1*-knockout mouse. (**A**) The remaining kidneys of wild-type (+/+) and *Cdcp1-*homozygous knockout (–/–) mice after UNX were subjected to immunofluorescence microscopy analysis using a primary antibody against CDCP1 pY734 and an Alexa Fluor 594-conjugated secondary antibody (magenta). (**B–D**) The remaining kidneys of wild-type (*Cdcp1* +/+) mice after UNX were subjected to immunofluorescence microscopy analysis using specific primary antibodies against Met pY1234-1235 (**B**), STAT3 pY705 (**C**), and Src pY418 (**D**) and an Alexa Fluor 488-conjugated secondary antibody (green). The endosome marker EEA1 was visualized using an Alexa Fluor 594-conjugated secondary antibody (magenta). (**E**) The remaining kidneys of *Cdcp1* wild-type (+/+) and *Cdcp1*-homozygous knockout (–/–) mice after UNX were subjected to immunofluorescence microscopy analysis using a primary antibody against MMP2 and an Alexa Fluor 594-conjugated secondary antibody (magenta). Proximal tubules were visualized by staining with FITC-LTL (green). The scale bars indicate 50 µm. (**F**) Schematic diagram summarizing differences in cellular events observed during compensatory renal growth of wild-type (+/+) and *Cdcp1*-homozygous knockout (–/–) mice.

**Fig. S9.**
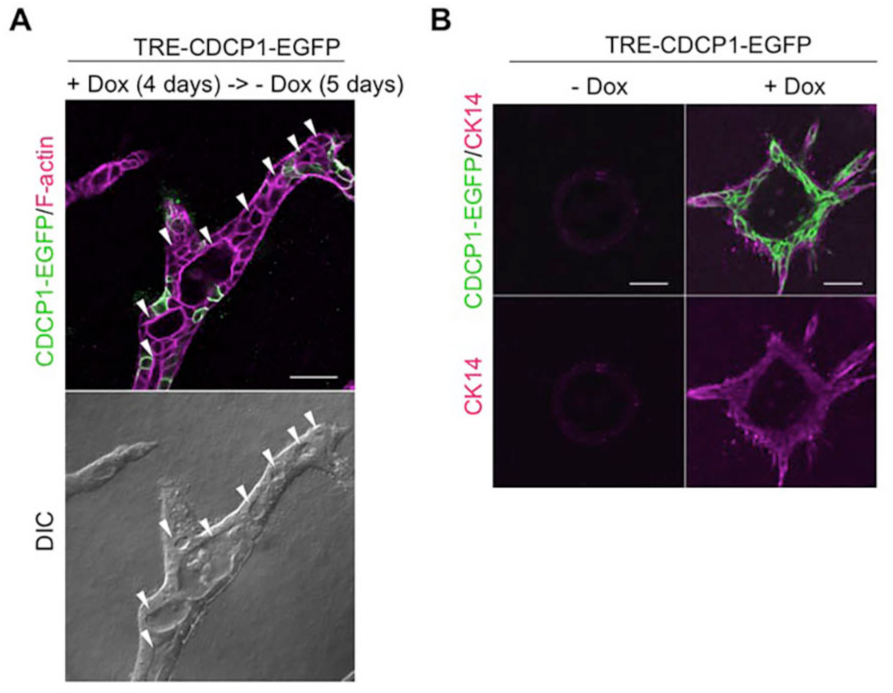
Effects of CDCP1 expression on the morphology of MDCK cysts. (**A**) CDCP1 downregulation induced the formation of a luminal structure. CDCP1–EGFP cysts embedded within the collagen matrix were incubated in the presence of Dox (1 µg/ml) for 4 days. Cysts were then incubated for an additional 5 days in the absence of Dox. (**B**) CDCP1 upregulation induced Cytokeratin 14 (CK14) expression. TRE–CDCP1–EGFP-harboring cysts were incubated with Dox (1 µg/ml) for 4 days. CK14 was visualized with a specific antibody (magenta). The scale bars indicate 50 µm.

**Fig. S10.**
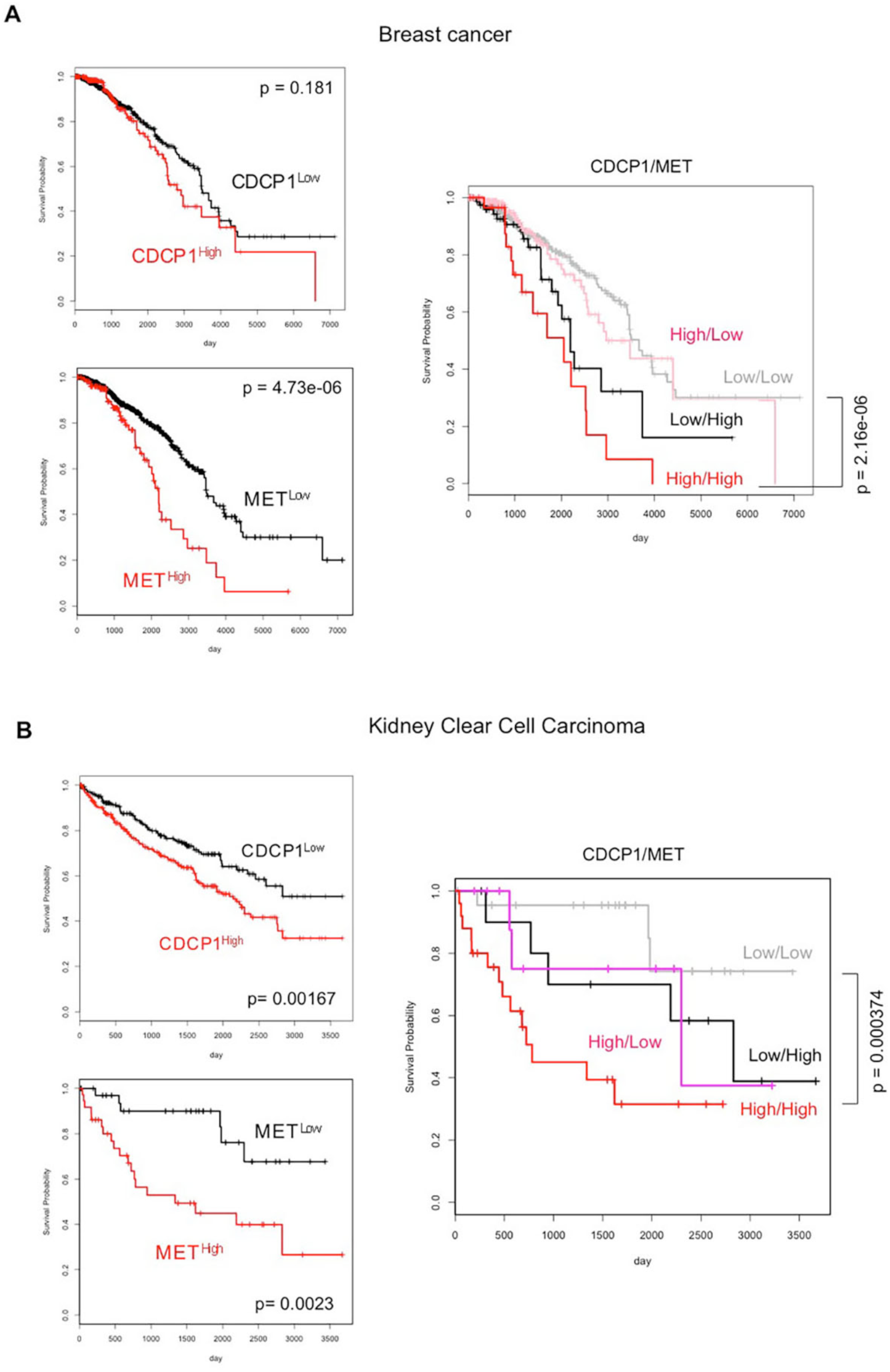
Co-upregulation of CDCP1 and Met correlates with poor prognosis in patients with breast and kidney cancer. (**A, B**) Correlations of CDCP1- and Met-expression levels with the prognoses of patients with breast cancer (**A**) or kidney clear cell carcinoma (**B**) were estimated using the Kaplan–Meier method, based on the transcriptome dataset from the TCGA project. Statistical significance was calculated by performing a log-rank test.

### Supplementary Tables

**Table S1.**
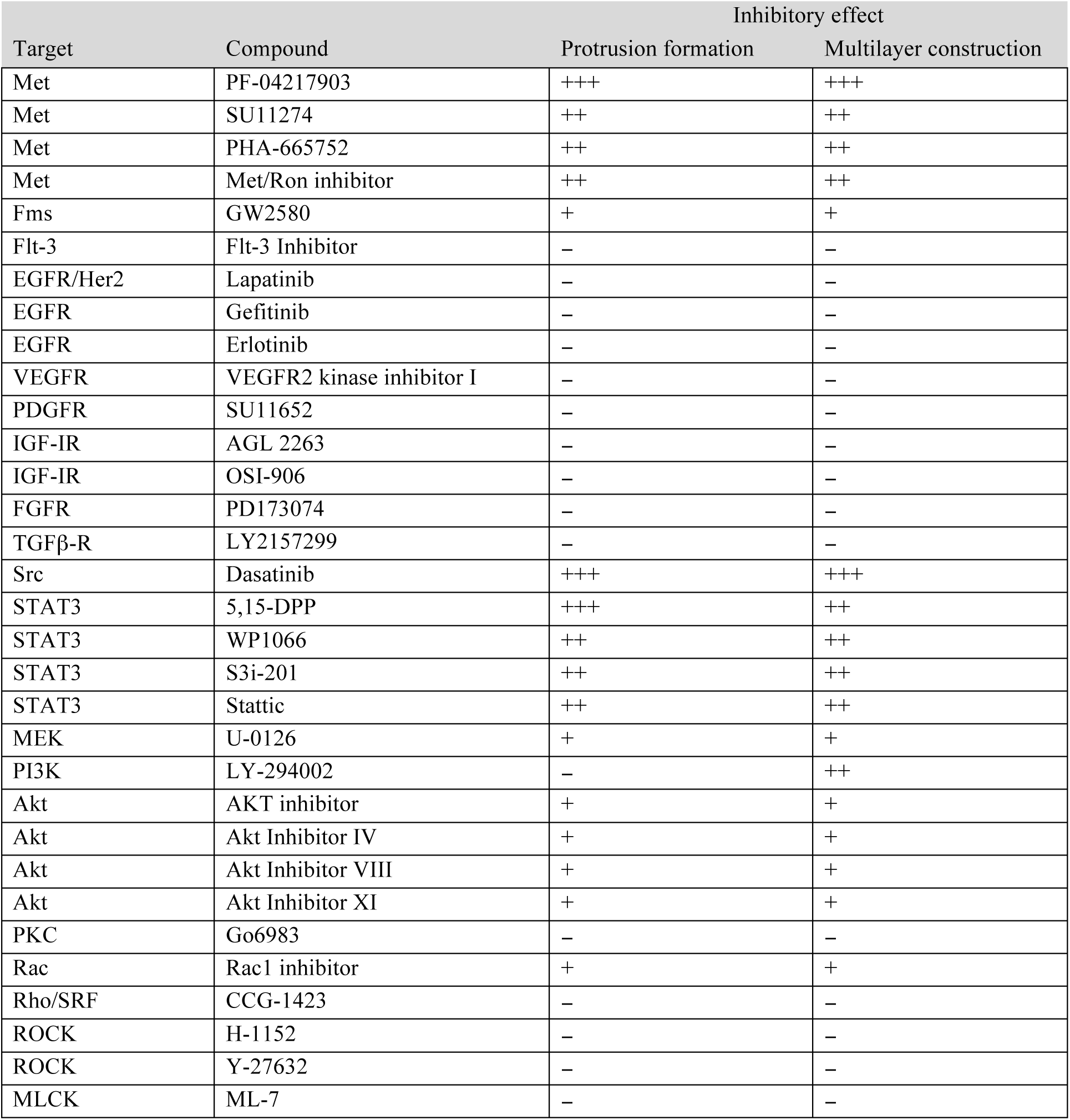
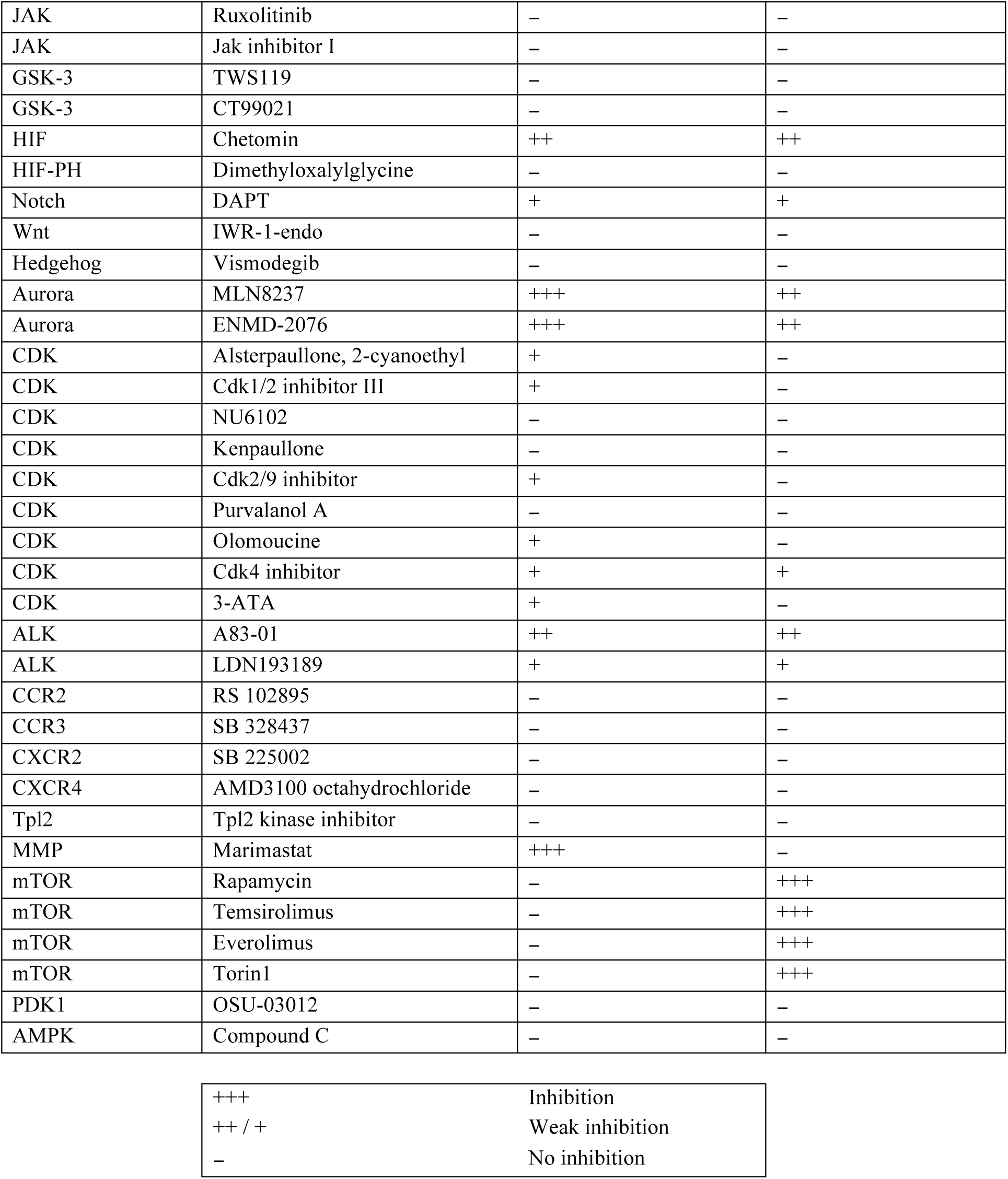
(1). Inhibitor screening for CDCP1-induced phenotypic changes in MDCK cysts
(2). Inhibitor screening for CDCP1-induced phenotypic changes in MDCK cysts

| Target | Compound | Inhibitory effect |  |
| --- | --- | --- | --- |
|  |  | Protrusion formation | Multilayer construction |
| Met | PF-04217903 | +++ | +++ |
| Met | SU11274 | ++ | ++ |
| Met | PHA-665752 | ++ | ++ |
| Met | Met/Ron inhibitor | ++ | ++ |
| Fms | GW2580 | + | + |
| Flt-3 | Flt-3 Inhibitor | – | – |
| EGFR/Her2 | Lapatinib | – | – |
| EGFR | Gefitinib | – | – |
| EGFR | Erlotinib | – | – |
| VEGFR | VEGFR2 kinase inhibitor I | – | – |
| PDGFR | SU11652 | – | – |
| IGF-IR | AGL 2263 | – | – |
| IGF-IR | OSI-906 | – | – |
| FGFR | PD173074 | – | – |
| TGFβ-R | LY2157299 | – | – |
| Src | Dasatinib | +++ | +++ |
| STAT3 | 5,15-DPP | +++ | ++ |
| STAT3 | WP1066 | ++ | ++ |
| STAT3 | S3i-201 | ++ | ++ |
| STAT3 | Stattic | ++ | ++ |
| MEK | U-0126 | + | + |
| PI3K | LY-294002 | – | ++ |
| Akt | AKT inhibitor | + | + |
| Akt | Akt Inhibitor IV | + | + |
| Akt | Akt Inhibitor VIII | + | + |
| Akt | Akt Inhibitor XI | + | + |
| PKC | Go6983 | – | – |
| Rac | Rac1 inhibitor | + | + |
| Rho/SRF | CCG-1423 | – | – |
| ROCK | H-1152 | – | – |
| ROCK | Y-27632 | – | – |
| MLCK | ML-7 | – | – |

|  |  |
| --- | --- |
| +++ | Inhibition |
| ++ / + | Weak inhibition |
| – | No inhibition |

**Table S1 (2).** Inhibitor screening for CDCP1-induced phenotypic changes in MDCK cysts
| Target | Compound | Inhibiting effect |  |
| --- | --- | --- | --- |
|  |  | Protrusion formation | Multilayer construction |
| JAK | Ruxolitinib | – | – |
| JAK | Jak inhibitor I | – | – |
| GSK-3 | TWS119 | – | – |
| GSK-3 | CT99021 | – | – |
| HIF | Chetomin | ++ | ++ |
| HIF-PH | Dimethyloxalylglycine | – | – |
| Notch | DAPT | + | + |
| Wnt | IWR-1-endo | – | – |
| Hedgehog | Vismodegib | – | – |
| Aurora | MLN8237 | +++ | ++ |
| Aurora | ENMD-2076 | +++ | ++ |
| CDK | Alsterpaullone, 2-cyanoethyl | + | – |
| CDK | Cdk1/2 inhibitor III | + | – |
| CDK | NU6102 | – | – |
| CDK | Kenpaullone | – | – |
| CDK | Cdk2/9 inhibitor | + | – |
| CDK | Purvalanol A | – | – |
| CDK | Olomoucine | + | – |
| CDK | Cdk4 inhibitor | + | + |
| CDK | 3-ATA | + | – |
| ALK | A83-01 | ++ | ++ |
| ALK | LDN193189 | + | + |
| CCR2 | RS 102895 | – | – |
| CCR3 | SB 328437 | – | – |
| CXCR2 | SB 225002 | – | – |
| CXCR4 | AMD3100 octahydrochloride | – | – |
| Tpl2 | Tpl2 kinase inhibitor | – | – |
| MMP | Marimastat | +++ | – |
| mTOR | Rapamycin | – | +++ |
| mTOR | Temsirolimus | – | +++ |
| mTOR | Everolimus | – | +++ |
| mTOR | Torin1 | – | +++ |
| PDK1 | OSU-03012 | – | – |
| AMPK | Compound C | – | – |

**Table S2.**
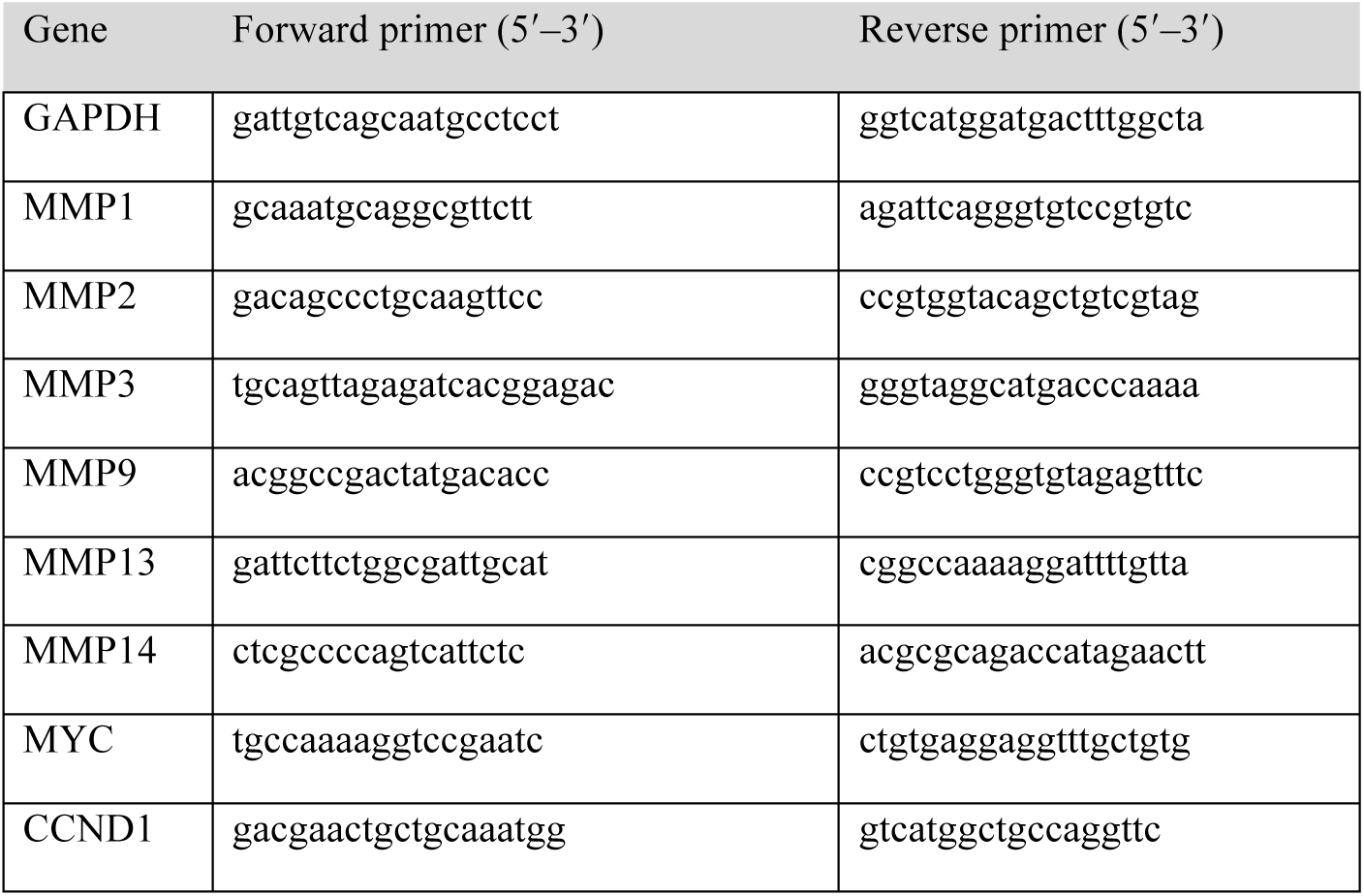
Sequences of primers used for quantitative RT-PCR.

**Table S3.**
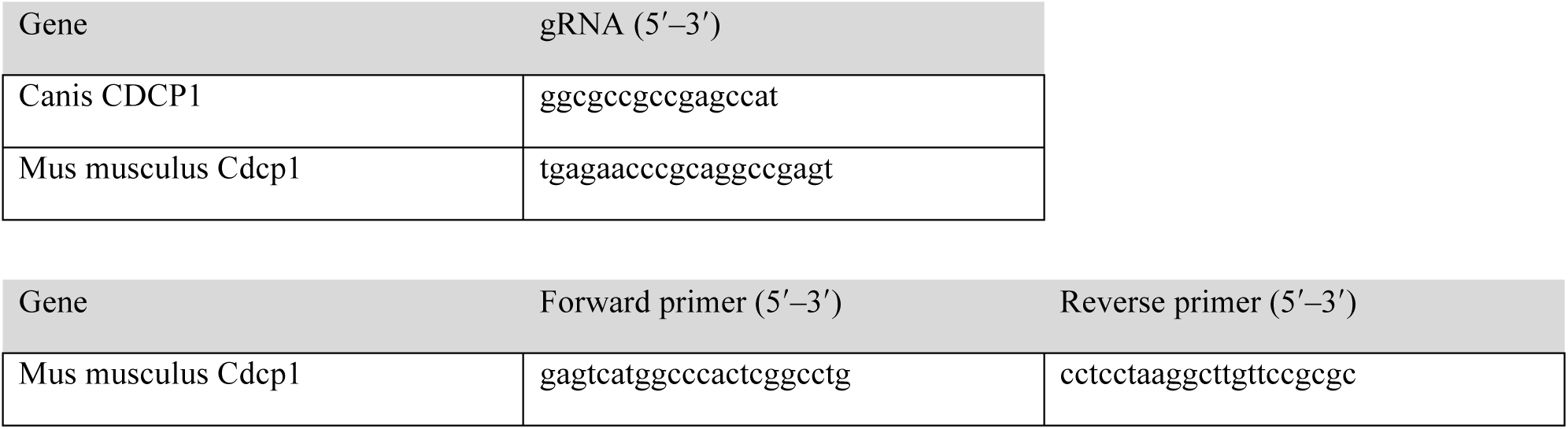
Sequences of gRNAs and genotyping primers.

**Supplementary Movie S1.** CDCP1-induced cell protrusion. CDCP1–EGFP MDCK cysts harboring mCherry-CAAX embedded within collagen matrix were incubated in the presence of Dox (1 µg/ml). Time-lapse images were captured at 30-min intervals for 40 h.

**Supplementary Data file S1.** Upstream regulator analysis using Ingenuity Pathway Analysis software.

